# Maternal effects on early-life gut microbiome maturation in a wild nonhuman primate

**DOI:** 10.1101/2021.11.06.467515

**Authors:** Alice Baniel, Lauren Petrullo, Arianne Mercer, Laurie Reitsema, Sierra Sams, Jacinta C. Beehner, Thore J. Bergman, Noah Snyder-Mackler, Amy Lu

## Abstract

Early-life gut microbial colonization is an important process shaping host physiology, immunity and long-term health outcomes in humans and other animals. However, our understanding of this dynamic process remains poorly investigated in wild animals, where developmental mechanisms can be better understood within ecological and evolutionary relevant contexts. Using 16s rRNA amplicon sequencing on 525 fecal samples from a large cohort of infant and juvenile geladas (*Theropithecus gelada*), we characterized gut microbiome maturation during the first three years of life and assessed the role of maternal effects in shaping offspring microbiome assembly. Microbial diversity increased rapidly in the first months of life, followed by more gradual changes until weaning. As expected, changes in gut microbiome composition and function with increasing age reflected progressive dietary transitions: in early infancy when infants rely heavily on their mother’s milk, microbes that facilitate milk glycans and lactose utilization dominated, while later in development as graminoids are progressively introduced into the diet, microbes that metabolize plant complex polysaccharides became dominant. Furthermore, the microbial community of nursing infants born to first-time (primiparous) mothers was more “milk-oriented” compared to similarly-aged infants born to experienced (multiparous) mothers. Comparisons of matched mother-offspring fecal samples to random dyads did not support vertical transmission as a conduit for these maternal effects, which instead could be explained by slower phenotypic development (and associated slower gut microbiome maturation) in infants born to first-time mothers. Together, our findings highlight the dynamic nature of gut colonization in early life and the role of maternal effects in modulating this trajectory in a wild primate.

## INTRODUCTION

The colonization of the gastrointestinal tract begins at birth and develops into a trajectory that can be highly variable between individuals [1–8]. Variation in the source and timing of postnatal microbial colonization influences somatic growth [9–12], neuroendocrine [13,14] and immune physiology [15–17], with health and fitness consequences that can extend across the life course [16,18,19]. In humans, for instance, infants that take antibiotics during the first year of life are more likely to develop allergies, asthma, and inflammatory bowel disease during childhood [20–23]. Germ-free rodent models demonstrate that at least some of these effects are causally related to the microbiome and are long-lasting. For example, germ-free rodents develop structural abnormalities of the gastrointestinal tract [15,24] that translate into immune system dysfunction later in life [25,26], an outcome that can only be partly reversed by introducing microbes during critical periods of development [27,28]. Despite the critical role that early-life gut microbial colonization plays in host development, research thus far has mainly focused on clinical studies in humans [14,22,29–31], complemented by experimental studies on laboratory rodents [32–35] and domestic animals [9,36–39]. Studies of wild animals are needed if we want to understand host-microbiome coevolution within a broader ecological and evolutionary context and without the confounding factors associated with medical interventions (e.g., Cesarean section, antibiotic use, formula feeding) [17,40,41].

The maternal microbiota drives gut microbiome assembly in offspring via vertical transmission of a large number of microbial lineages [42–46]. Vertical transmission is thought to be particularly strong in mammals due to viviparity and extended periods of lactation and post-weaning maternal care [47]. The first important exposure to microbes occurs during birth, when infants are inoculated with maternal vaginal, fecal, and skin microbiota [3,5,8,29,44,48]. Postnatally, vertical transmission is primarily accomplished through nursing, with numerous microbes and milk glycans (i.e., oligosaccharides) transmitted through milk that, together, determine the microbial composition of the infant’s gut [42,49,50]. While milk microbes directly seed the offspring’s gut, milk glycans promote the growth of beneficial microbes, such as *Bifidobacterium* and *Bacteroides*, that in turn break the glycans down into forms usable by the host [51–53]. Although breastmilk is the most obvious route by which vertical transmission takes place [10,49,50,54], studies on humans suggest that the maternal gut microbiome is also a major source of transmitted strains [42,44,55,56]. Maternal gut microbes might be transmitted to offspring via milk, as the gastrointestinal tract is hypothesized to be the major reservoir of microbes colonizing the mammary gland (the enteromammary pathway) [49,57]. Alternatively or additionally, mothers may transmit gut microbes to offspring via preferential body contact [58], a mechanism that suggests vertical transmission can continue in some capacity past weaning [47]. Because maternal microbial taxa are the first to colonize and tend to be better adapted to the gut ecological niche compared to other environmental microbes, they often persist longer in offspring than those acquired from other sources [42,56,59].

Recent studies suggest that microbiome-mediated maternal effects are indeed possible. In several mammals, maternal traits, such as parity (i.e., the number of times a mother has given birth), have been associated with differences in the composition of both maternal [39,60] and offspring microbial communities [10,39]. In nonhuman primates (vervet monkeys: *Chlorocebus pygerythrus*), infants born to low-parity mothers harbored reduced microbial diversity and a greater abundance of *Bacteroides fragilis* [10], a bacterium derived from the milk microbiota that is specialized in digesting milk glycans [61,62]. In turn, infants from low-parity females grew faster, suggesting that low-parity mothers may compensate for poor milk production by vertically transmitting milk microbes that could help infants extract more energy from lower milk volumes [10]. Such strategies may be broadly beneficial to dyads in which mothers cannot provide adequate nutritional resources to offspring (e.g., low-ranking mothers) [63,64]. Thus, maternal vertical transmission of microbes may be an important mechanism of phenotypic plasticity during lactation [64,65].

Primates are particularly relevant models for understanding postnatal microbiome development and maternal effects because they are closely related to humans, display prolonged lactation periods, and engage in high maternal investment [66,67]. Furthermore, maternal condition (e.g., energetic status) and maternal traits (e.g., dominance rank, social integration parity) are known to influence offspring developmental and long-term fitness outcomes [68–71]. Although studies on host-associated microbial communities in wild primates are emerging, many remain limited in scope, hampered by cross-sectional samples and small sample sizes of unweaned infants (particularly in the first few weeks of life), which together prevent longitudinal characterization of gut microbial colonization processes [72–75]. Here, we used dense cross-sectional and longitudinal monitoring to characterize gut microbial colonization during the first three years of life and assess the role of maternal effects in shaping offspring maturation trajectories in wild gelada monkeys (*Theropithecus gelada*). Geladas live in the high-altitude plateaus of Ethiopia and have a specialized graminivorous diet (at times, comprising 90% grass) [76,77], which strongly shapes the composition and function of the adult gut microbiome [78,79]. Because geladas live in polygynous reproductive units that range together in larger bands (comprised of 200 or more individuals) [80], we are able to monitor over 50 immatures at any given time, offering an unprecedented sample size to examine gut microbial characteristics during early life in a wild primate. We used 16s rRNA amplicon sequencing on 525 fecal samples from 89 immatures to profile changes in gut microbiome diversity, composition, and function during the first three years of life (N=5.9±5.5 samples per individual, range:1-18, **Figure S1**). In our population, geladas reach weaning at approximately 1.5 years of age and become sexually mature around 4.6 years [81]; and maternal characteristics, such as parity and dominance rank, are known to influence inter-individual variation at both of these developmental milestones [Feder et al., in revision; Lu et al., unpublished]. We predicted that early life microbial changes would reflect dietary transitions associated with weaning, as infants transition from milk to a plant-based diet [5,48,82,83]. We also predicted that maternal traits, such as dominance rank and parity would be associated with inter-individual differences in gut microbiome diversity, composition, and function in offspring. More specifically, we predicted that infants born to primiparous and low-ranking mothers would have a microbiome more functionally adapted to digest milk to compensate for poorer maternal energetic allocation during lactation. Lastly, we tested if we could detect evidence of vertical transmission between mother and offspring using fecal-fecal comparisons of mother-infant dyads (with 398 matched fecal samples between mother and offspring collected on the same day throughout development) and if greater vertical transmission in certain females (e.g., low rank, first-time mothers) could be the conduit for putative maternal effects on offspring’s microbiome composition. We expected a stronger signal of vertical transmission in early life [10,42,44,55,56], likely driven by a combination of greater microbial transfer via milk when infants are nursing and are also in more frequent body contact with their mother.

## RESULTS

### General pattern of gut microbiome maturation in geladas

We characterized the gut microbiome across 525 immature gut microbiome samples, and detected 3,784 Amplicon Sequence Variants (ASVs) (mean±SD per sample: 728±261, min-max: 65-1,498) belonging to 19 phyla and 76 families. The gut microbiome composition of immature geladas changed quickly following birth, with an initial phase of taxonomic succession and diversification during the first few months of life, followed by a progressive stabilization of the overall community (Figures 1A,B).

**Figure 1.**
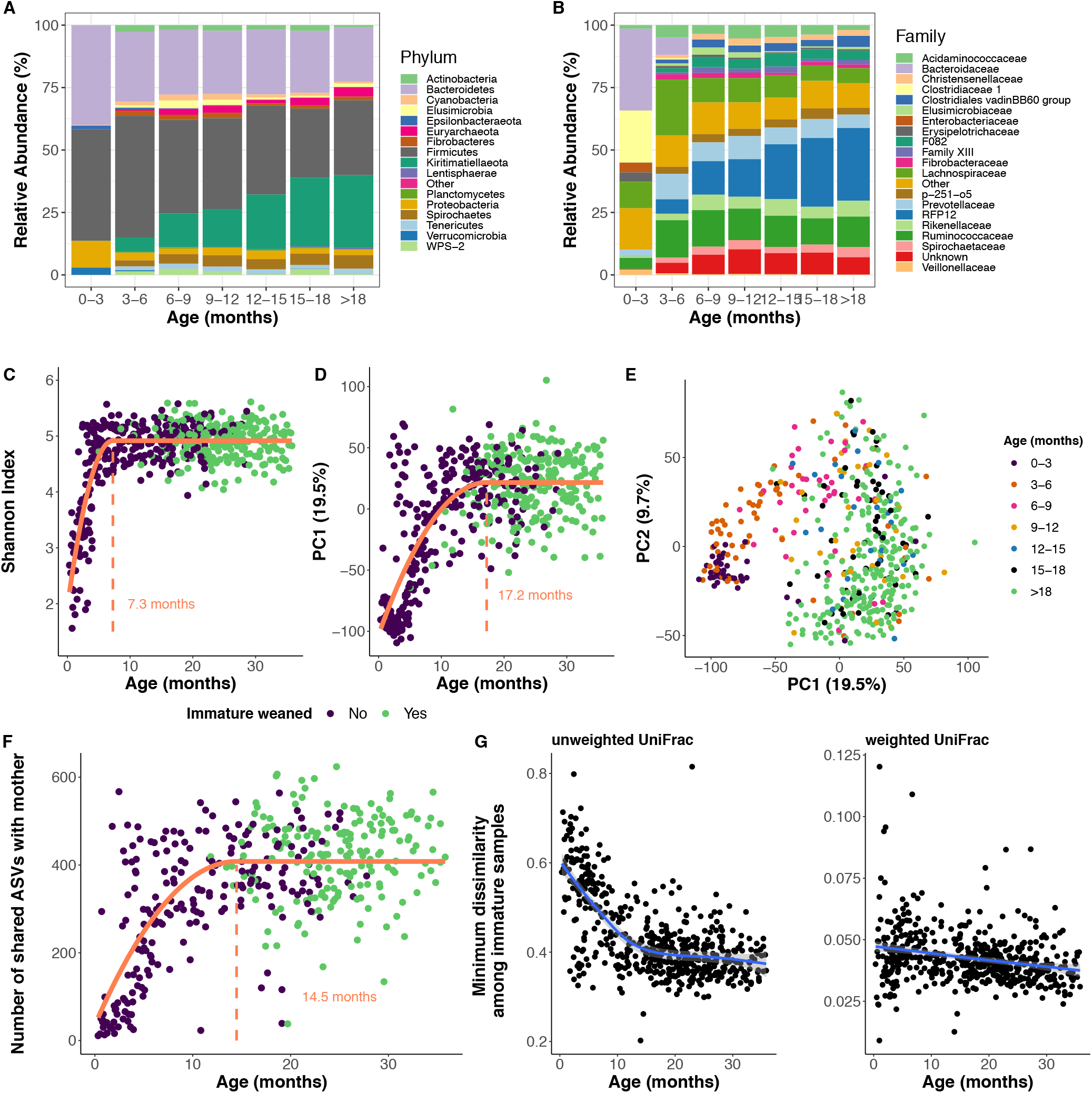
Gut microbiome taxonomic assembly in the first three years of life in immature geladas. (A, B) Taxonomic composition of the immature gelada gut microbiome at the phylum and family level as a function of age. The relative abundance of each taxon was calculated per sample by dividing the counts of the taxa by sequencing depth, and then averaged across samples belonging to the age category of interest. Age was split into categories for visualization purposes, but analyses treated age as a continuous variable. (C) Age-associated pattern of alpha diversity within samples, as calculated by the Shannon index (richness and evenness of Amplicon Sequencing Variants, ASVs). The vertical line represents the critical point of inflexion (calculated using quadratic plateau models) representing the age at which alpha diversity converges to adult-like patterns. The dataset was rarefied at 20,000 reads for the figure. (D,E) Age-associated pattern of beta diversity. A Principal Component Analysis (PCA) was used to ordinate the samples based on the Aitchison dissimilarity index (which is simply the Euclidean distance after centered-log-ratio transformation of the raw counts). Panel D represents the projection of the first principal component (PC1) that is best explained by the age of immatures. The vertical line represents the critical point of inflexion (calculated using quadratic plateau models) representing the age at which beta diversity converges to adult-like patterns. Panel E shows how age structures the gut microbiome composition of immatures on the two first principal components. (F) Comparison of gut microbiome composition between mother and offspring, as assessed using 398 matched mother-infant pairs of fecal samples collected on the same day. Here, the number of shared ASVs between pairs of samples is represented. The vertical line represents the critical point of inflexion (calculated using quadratic plateau models) representing the age at which the number of shared ASVs stabilizes to its maximal value. The dataset was rarefied at 20,000 reads for the calculation. (G) Age distribution inter-individual variability in gut microbiomes using the ASV-level unweighted and weighted UniFrac distances. This score was calculated as the minimum pairwise dissimilarity value from a beta diversity matrix for a given immature sample, and captures how dissimilar a sample is from its nearest neighbor, given all other gut microbiome samples in the immature cohort. Higher values indicate a more distinct gut microbiome from the population. The dataset was rarefied at 20,000 reads for the calculation.

To characterize broad changes in gut microbial community composition across development, we first focused on patterns of alpha diversity (i.e., the microbial diversity within a sample) and beta diversity (i.e., the overall difference of composition between samples). The Shannon Index of alpha diversity was initially low in early life and increased rapidly with age (GAMM: edf=7.2, *P*<2.0×10^-16^) (Figure 1C**, Table S1**), converging to adult-like values at 7.3 months (nonlinear quadratic plateau model: R^2^=0.62) (see **Figure S2, Table S1** for similar results on alternative alpha diversity metrics). Furthermore, age was one of the strongest predictors of the difference in microbial composition between samples (PERMANOVA based on Aitchison dissimilarity metric of beta diversity: R^2^=0.75, *P*<9.9×10^-05^, **Table S2**, Figures 1D,E) and samples clustered tightly by age on the first axis (PC1) of a Principal component analysis of beta diversity (Pearson correlation coefficient between age and PC1=0.62, *P*<2.2×10^-16^). Compared to alpha diversity, beta diversity reached an adult-like composition later in development, at 17.2 months (nonlinear quadratic plateau model between PC1 and age: R^2^=0.55; Figure 1D), which is approximately the age at which gelada mothers return from lactational amenorrhea and resume reproductive cycles [81]. Other important structuring factors of the immature gut microbiome included infant identity (R^2^=0.24) and group membership (R^2^=0.05) (**Table S2**).

To assess the compositional maturation of the gut microbiome of immature geladas relative to the maternal gut microbiome across age, we calculated the number of shared ASVs and beta diversity dissimilarity (unweighted and weighted UniFrac) between 398 matched immature-mother pairs of fecal samples collected the same day. As offspring got older, they shared an increasing number of bacteria with their mother (GAMM: effective degree of freedom, edf=4.7, *P*<2.0×10^-16^; **Table S3**, Figure 1F) and became more similar to maternal (i.e., adult-like) gut microbiome composition (unweighted UniFrac: edf=4.7, *P*<2.0×10^-16^; weighted UniFrac: edf=3.5, *P*<2.0×10^-16^; **Table S3, Figure S3**). Convergence with maternal gut occurred at 14.5 months for the number of shared ASVs (nonlinear quadratic plateau model: R^2^=0.44; Figure 1F) and 14.8-15.5 months for beta diversity dissimilarity (R^2^=0.48 for unweighted UniFrac and R^2^=0.17 for weighted UniFrac; **Figure S3**).

Despite the strong age-related patterns noted above, inter-individual variability in composition (as measured by the minimal pairwise beta diversity dissimilarity value among immature samples, see Methods) was higher among younger infants compared to older juveniles (Figure 1G). Some young infants (∼3-6 months) in particular had a gut microbiome that were relatively mature (i.e. adult-like) for their age (Figure 1D). Such “individuality” in the gut microbiome in early life was likely driven by the presence of rare taxa, since the pattern was stronger using unweighted UniFrac (which does not take into account taxa abundance) as opposed to weighted UniFrac measures of beta diversity (Figure 1G).

### Taxonomic and functional changes during development

To characterize age-associated changes in microbial composition and function, we used autoregressive integrated moving average (ARIMA) models to identify significant developmental changes in the abundance of each microbial taxa (at the family and genus levels) and each predicted functional pathway (at the metabolic level: KEGG Orthologs, KOs and enzymatic level: Enzyme Commission numbers, EC [84]). We then used hierarchical clustering to group microbial taxa and functional pathways based on similar age-related abundance trajectories. Maturational trajectories fell into one of four distinct clusters at both the taxonomic (Figure 2**, S4; Table S4**) and functional (KOs: **Figure S5-S6, Table S5**; EC: **Figure S7, Table S6**) levels.

**Figure 2.**
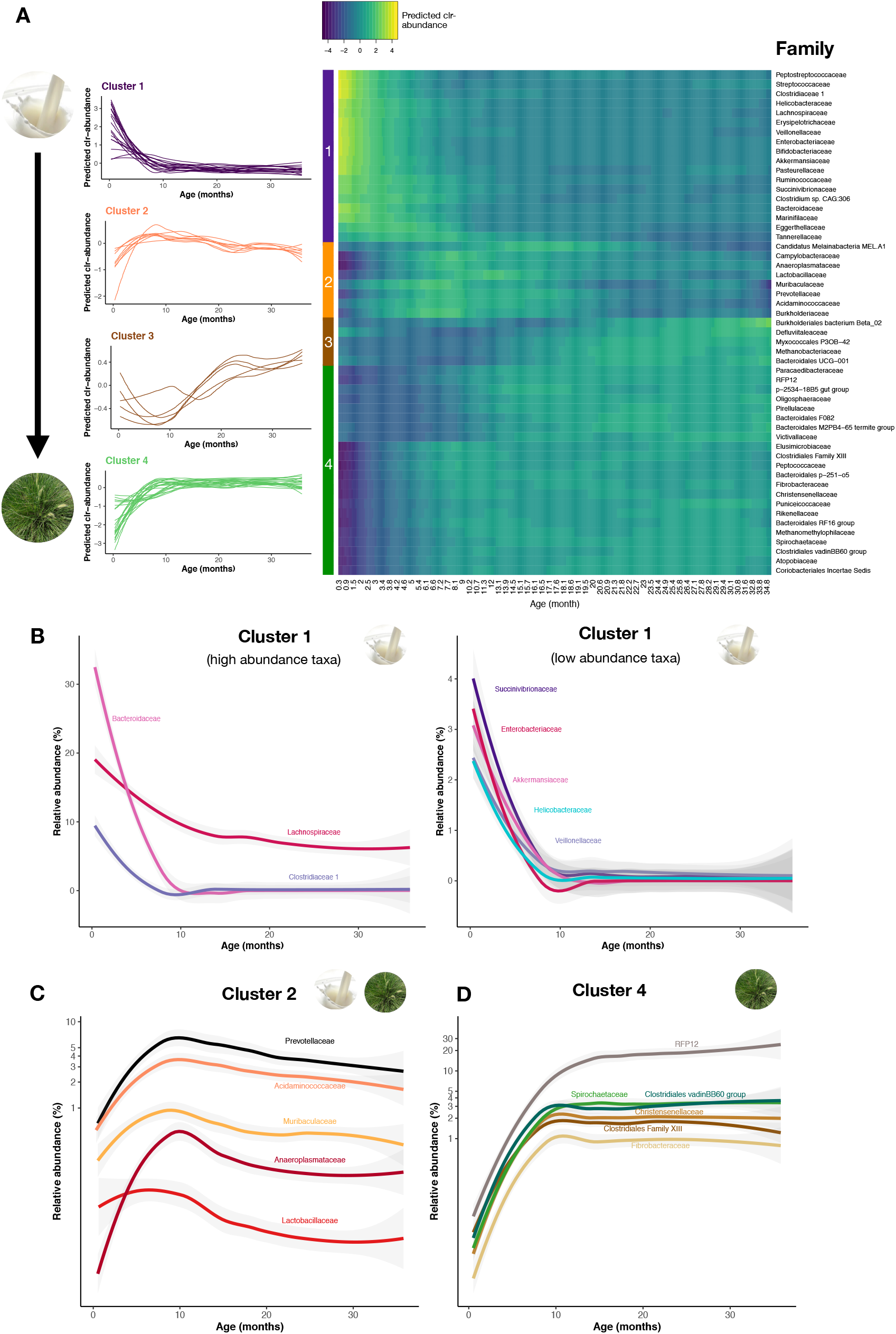
Age-associated changes in microbial composition at the family level. (A) Heatmap of the microbial families exhibiting a significant chronological trend as a function of age (fitted values from ARIMA models and predicted using LOESS regression per taxa as a function of age). Values represent z-score normalized counts after centered-log-ratio (clr) transformation. Hierarchical clustering was used to group age-dependent trajectories into four clusters exhibiting similar chronological trends. Color bars on the left side identify the clusters. Taxa (i.e. rows) are ordinated on the heatmap using correlation as distance function. All microbial families above 0.01% abundance were analyzed (N=55) and the 53 that displayed a significant trend are represented. (B) Relative abundance of 8 functionally important microbial families that are enriched in early life (belonging to cluster 1), as a function of age. The age-dependent trajectories were calculated on clr-transformed counts, but here for interpretation purposes we represent the LOESS regression on the raw relative abundance across samples (same for panels C and D). (C) Relative abundance of 5 functionally important microbial families that peak in abundance around 10 months of age (belonging to cluster 2). Relative abundance is represented on a log-scale to accommodate high and low abundance families together. (D) Relative abundance of 8 functionally important microbial families that are enriched in later life (belonging to cluster 4). Relative abundance is represented on a log-scale to accommodate high and low abundance families together.

#### Cluster 1: The early-life microbiome is adapted to process milk

Cluster 1 contained microbes that were abundant during the earliest months of infancy (18 families: Figure 2A**, Table S4** and 39 genera: **Figure S4, Table S4**) and are broadly involved in using and fermenting milk sugars (see supplemental results 1 for additional details on cluster 1). These early colonizers comprised bacteria that break down milk glycans (Bacteroidaceae, Bifidobacteriaceae) and lactose (Streptococcaceae, [*Ruminococcus] gnavus group*) and other groups that ferment glycans and lactose into butyrate (Lachnospiraceae: *Lachnoclostridium*, *Blautia*, *Anaerostipes*, and Ruminococcaceae: *Faecalibacterium, Butyricicoccus, Butyricimonas*) or propionate (Veillonellaceae) (Figures 2B **and** 3A). *Bacteroides* appeared to be the main degrader of milk glycans in geladas, representing the most abundant genus in early life (∼30% of the gut microbes at 1 month) (Figure 3A). One *Bacteroides* ASV – *B. fragilis,* a proficient degrader of milk glycans [61] – was particularly abundant in early life (i.e., with a high loading score on PC1, **Table S7**). By contrast, *Bifidobacterium* – an important milk glycan degrader in humans – was present at extremely low abundance across development (<0.01% at 1 month in geladas vs ∼40% in humans [85]) (Figure 3A).

**Figure 3.**
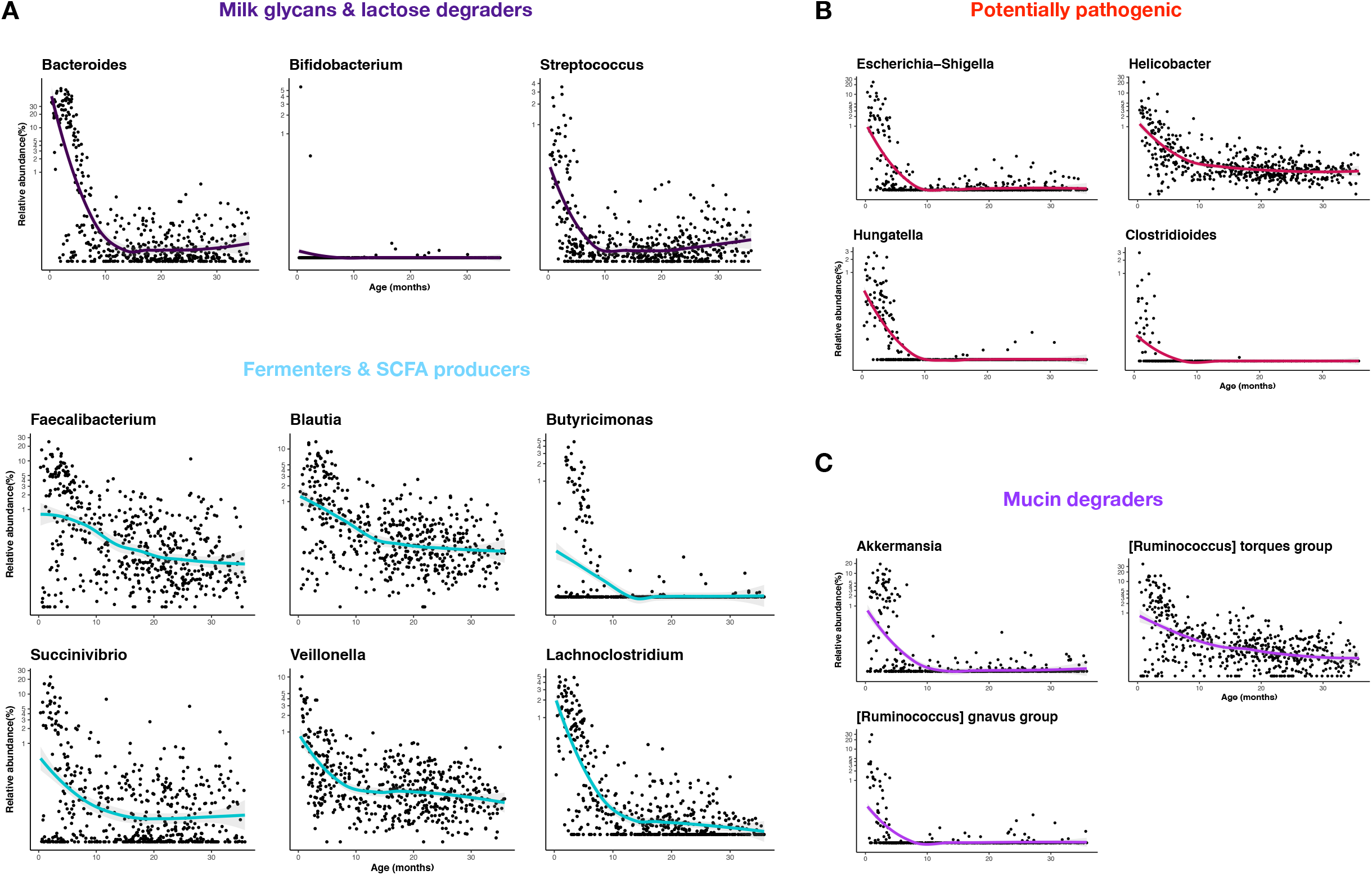
Composition of the early-life microbiota at the genus level. Relative abundance of functionally important genera in early life, as a function of age. The age-dependent trajectories were calculated on clr-transformed counts. For visualization purposes however, we represent the LOESS regression on the raw relative abundance across samples (on a log-transformed scale).

Functional cluster 1 also reflected the involvement of the gut microbiome in milk utilization and immunity pathways (metabolic cluster 1: **Figures S5-S6, Tables S5** and enzymatic cluster 1: **Figure S7, Table S6**). Young infant gut microbiomes contained high levels of bacterial genes involved in carbohydrate metabolism, notably in the catabolism of fructose, mannose, and galactose (3 abundant milk sugars [86]), and in the conversion of sugars to energy (e.g., via glycolysis/gluconeogenesis, pyruvate metabolism, pentose phosphate pathway) (Figure 4A). Similar functional signatures of cluster 1 were also apparent at the enzymatic level, as the gut microbes encoded a specialized enzymatic toolkit (alpha and beta glucosidase, alpha and beta galactosidase, fucosidase, sialidase, beta-hexosaminidase) necessary to cleave complex milk glycans (**Figure S8, Table S6**). *Bacteroides* was the main microbial group encoding those enzymes (**Figure S8**), confirming its central role in milk glycan degradation in geladas.

**Figure 4.**
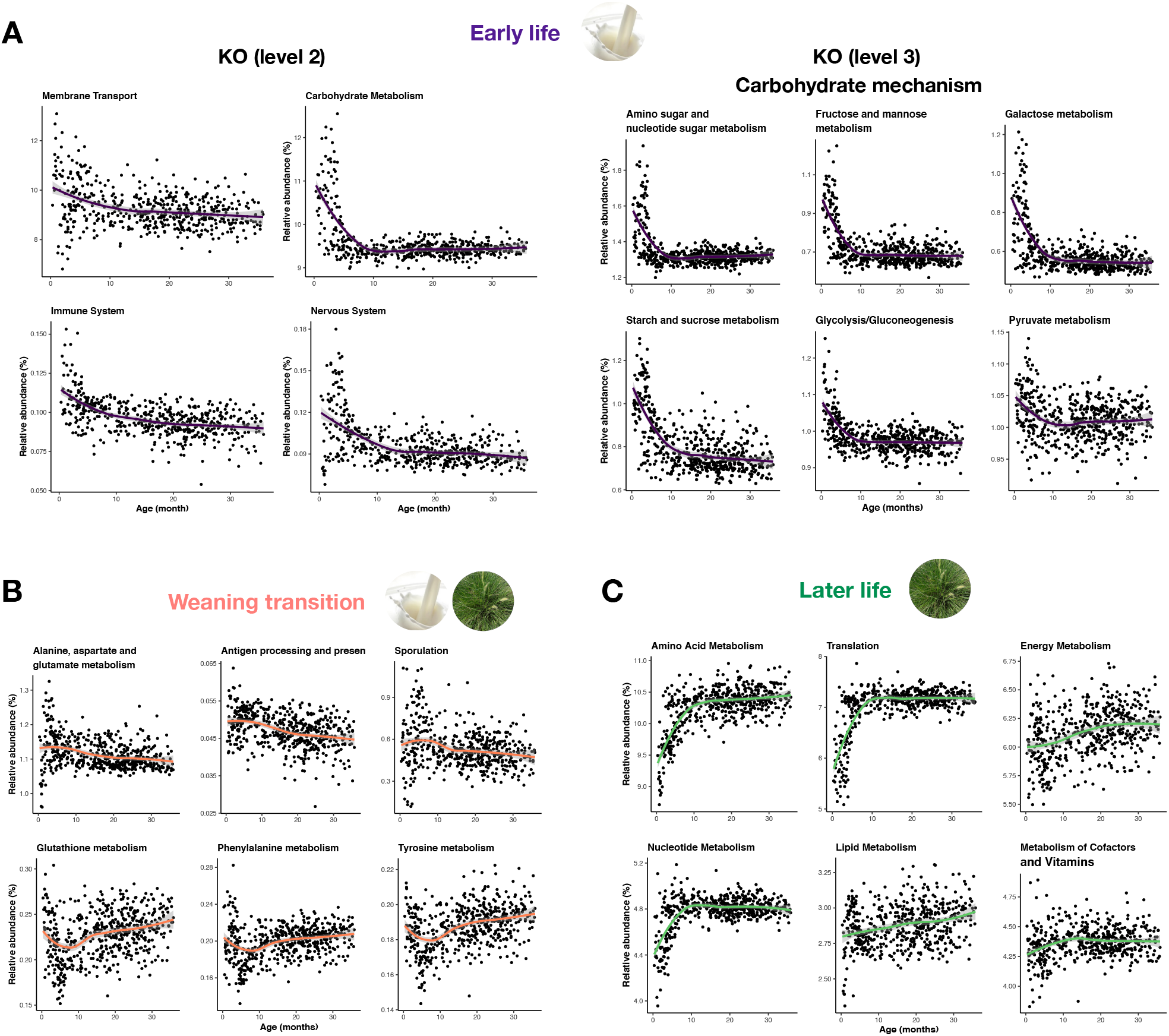
Age-associated changes in the functional profile of the gut microbiome of immatures based on predicted KEGG orthologs (KO) metagenomes. (A) Relative abundance of metabolic pathways (left: KO level 2 and right: KO level 3 of the carbohydrate metabolism) enriched in early life, as a function of age. (B) Relative abundance of metabolic pathways (KO level 3) that peak in abundance or decrease in abundance at 10 months of age. (C) Relative abundance of metabolic pathways (at KO level 2) enriched in later life, as a function of age. In all plots, the average curve is the LOESS regression on the raw relative abundance across samples.

Interestingly, cluster 1 also included several putatively pathogenic genera (Figure 3B), including some bacterial species most responsible for enteric infections and diarrheal diseases in human newborns and captive animals (e.g., *Clostridioides difficile*, *Helicobacter macacae*, *Clostridium butyricum, C. perfringens* [87–91]) (**Table S7**). It also included 3 major groups of mucin-degrading bacteria (*Akkermansia*, [*Ruminococcus] gnavus group* and [*Ruminococcus] torques*) (Figure 3C) that are involved in the development of the intestinal mucosa, a primary line of immune defense [92]. These taxa reflect the importance of the developing immune system at this stage in life. In line with this interpretation, the early life microbial metabolic pathways tended to be more involved in processes related to the host immune system (e.g NOD-like receptor) and nervous system (e.g., glutamatergic synapse pathway) (Figure 4A**, Tables S5**).

#### Clusters 2 & 3: The weaning transition is accompanied by important gut microbial rearrangements

Around 10 months of age, a small number of microbial taxa (Figures 2A **and S4, Tables S4**) and metabolic pathways (**Figures S5-S6, Tables S5**) peak (cluster 2) or decrease (cluster 3) in abundance. Of these changes, the most notable included peaks in Lactobacillaceae (genus *Lactobacillus*), Prevotellaceae, and Lachnospiraceae (cluster 2, Figure 2C). While Lactobacillaceae is a keystone lactic acid bacterial group producing large amounts of lactate from milk sugars, Prevotellaceae and Lachnospiraceae (Figure 2C) are fiber-degrading genera. These transient shifts highlight the role of the gut microbiome in digesting both milk and plant items at this age.

Taxonomic changes at ten months translated at the functional level into a remodeling of the metabolism of amino acids, with an increase in microbial genes involved in alanine, aspartate, glutamate, cysteine, and methionine metabolism (cluster 2), and a decrease in microbial genes involved in phenylalanine (found in breast milk), glutathione (antioxidant typically enriched in the first weeks of life in humans), and tyrosine metabolism (cluster 3) (Figure 4B**, Tables S5**). Microbial genes involved in sporulation and germination were also more highly expressed (Figure 4B**, Table S5**), suggesting some changes in persistence strategy from the spore-forming microbes in the gut.

#### Cluster 4: The later-life gut microbiome is adapted to a plant-based diet

Cluster 4 was characterized by 22 families (Figure 2A**, Table S4**) and 63 genera (**Figure S4, Table S4**) that increased sharply with age and plateaued in older immatures (from 10 months of age onward), including cellulolytic (Spirochaetaceae, Fibrobacteraceae, *Cellulosilyticum*) and fermentative taxa (Lachnospiraceae, Clostridiales Family XIII, several genera from Prevotellaceae and Ruminococcaceae), as well as RFP12 (Figure 2D), which are all commonly found in adult geladas [79]. These taxa are involved in metabolizing complex plant polysaccharides found in graminoid leaves and roots, which comprise the majority of the adult gelada diet.

At the functional level (cluster 4, **Figures S5-S6, Tables S5**), the gut of old immatures harbored more bacterial genes involved in energy, amino acid, and lipid metabolism and in the regulation of genetic expression and bacteria growth (nucleotide metabolism, replication and repair, genetic information processing and translation) (Figure 4C), a functional profile that is typical of the adult gelada gut [79].

### Maternal effects on offspring gut microbiome composition and function

We next examined whether inter-individual variability in gut microbiome composition early in life (Figure 1D,G) could be explained, in part, by maternal traits, including maternal dominance rank and parity. We ran these analyses using (i) all samples (0-3 years, N=525), but since we predicted that maternal effects on the offspring microbiome would be strongest in early life (when infants are still nursing), we also ran separate analyses only focusing on (ii) young infants (<12 months of age, still relying largely on milk, N=184) and (iii) old immatures (>18 months, relying largely on plants, N=259). Note that we ran separate analyses for each age group because it is not possible to fit an interaction between a smooth term (i.e., age) and covariates (i.e., maternal attributes) in GAMMs.

Maternal dominance rank did not influence the alpha or beta diversity (**Tables S1-S2, Figure S9**) of immature gut microbiomes, nor did it predict differences in microbial families, genera, or functional pathways (**Tables S8-S10** for (ii) young infants, results not shown for (i) all immatures or for (iii) old immatures). Maternal parity also did not exert a significant influence on the diversity, composition, or relative abundance of taxa in the immature gut microbiome (**Tables S1, S2, S8**). However, parity was significantly associated with the relative abundance of several microbial metabolic pathways (**Table S9**) and enzymes (**Table S10**) during the first 12 months of life (results non-significant for (i) all immatures or (iii) old immatures). Namely, infants born to primiparous females had functional profiles more typical of early life (<12 months) and related to milk digestion, both at the metabolic and enzymatic levels. Their gut microbes were more involved in carbohydrate metabolism (e.g., galactose, fructose and mannose metabolism), cellular processes and signaling, and nervous system function (Figure 5A); and they harbored a higher abundance of key enzymes that cleave milk glycans (**Figure S10**). By contrast, young infants (<12 months) born to multiparous females had a more functionally “mature” gut microbiome for their age, with higher abundance of later-life microbial pathways such as amino acid metabolism and nucleotide metabolism (Figure 5B). To determine why maternal parity had an effect at the functional level but not at the taxonomic level, we examined the bacterial taxa that showed a statistical trend to be more abundant in young infants (<12 months) born to primiparous females (i.e., with p-values<0.1 before FDR correction, **Table S9**). Offspring of primiparous mothers indeed tended to harbor a higher abundance of microbial taxa involved in milk digestion (e.g., Lachnospiraceae, Bacteroidaceae, Clostridiaceae 1) (**Figure S11**, see supplemental results 2), which suggests that individual taxa exert small additive effects that were only detected at the functional level.

**Figure 5.**
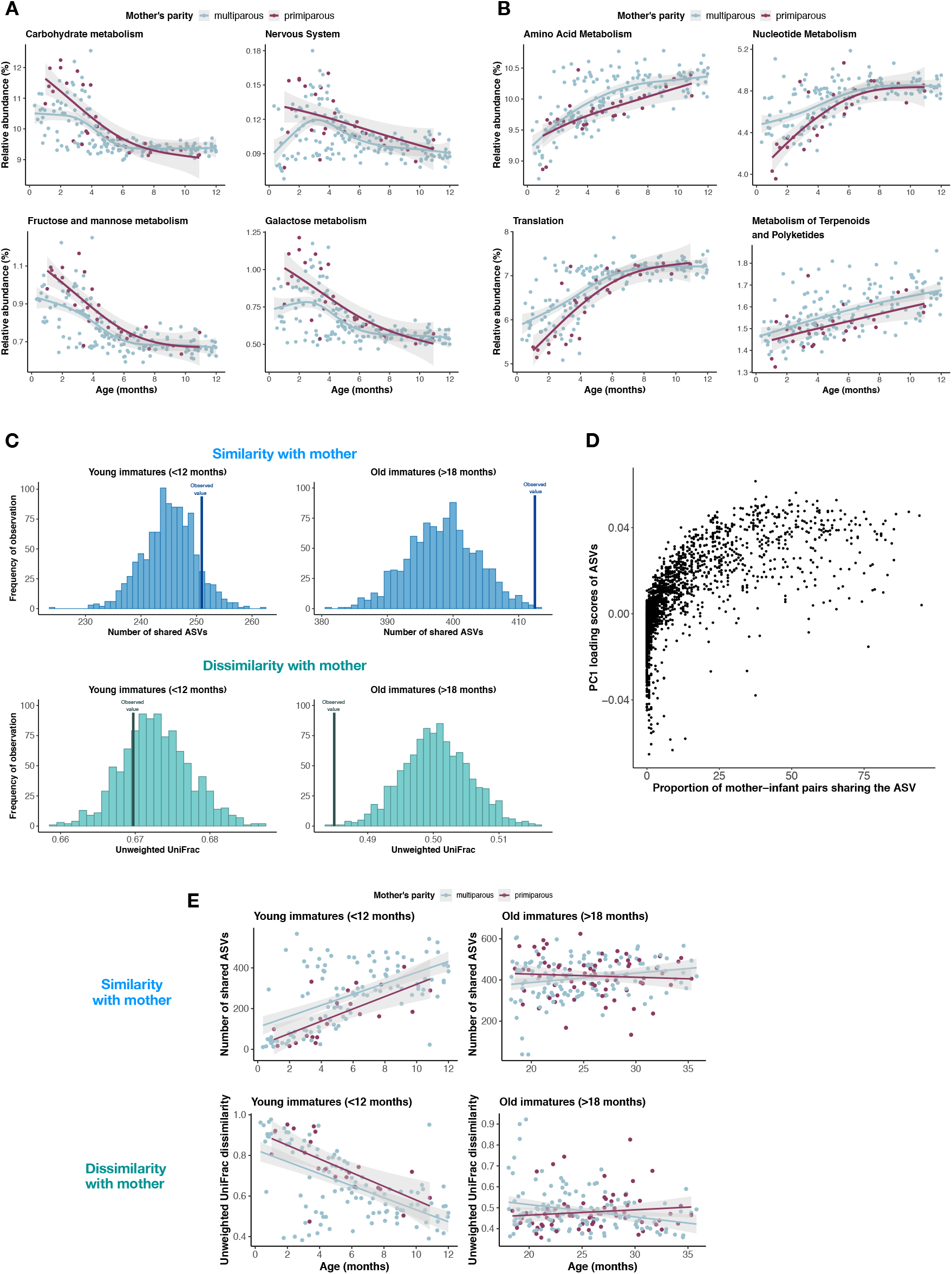
The effect of maternal parity on offspring’s gut microbiota functional capacity. (A) Metabolic pathways (KO level 2 for upper row and KO level 3 for lower row) that are more abundant in infants <12 months) born to primiparous females than infants of multiparous females. (B) Metabolic pathways (KO level 2) that are less abundant in infants (<12 months) born to primiparous females than infants of multiparous females. (C) Results of the nonparametric resampling approach testing if offspring share more Amplicon Sequencing Variants (ASVs) and have a more similar gut microbiome composition (unweighted UniFrac dissimilarity) to their mother than to random adult females of the population. The histograms show the random distribution of the metric of interest (i.e. when matching each infant sample to a random female sample collected during the same season, with 1000 repetitions). The vertical line shows the observed value of the metric (i.e. between the actual mother-offspring pairs of fecal samples collected the same day). This analysis was performed separately on young (nursing) infants (aged 0-12 months, N=136 samples) and old immatures (>18 months, N=201) that were likely weaned. (D) Composition of the shared ASVs between young infants (<12 months) and their mothers. The ASVs that are commonly shared between mother-offspring pairs (e.g. among > 70% of the pairs) in early life tend to have high loading scores of the first principal component (PC1) of a Principal Component Analysis ordination (based on Aitchison distance). Since PC1 strongly correlates positively with age, these shared ASVs are characteristic of later life. (E) Vertical transmission differs for primiparous and multiparous females in early life. Young infants (<12 months) born to primiparous females share fewer ASVs with their mother and have a more dissimilar gut microbiome composition (unweighted UniFrac dissimilarity) compared to their mother than offspring born to multiparous females. Later in life (>18 months), immatures born to primiparous and multiparous females are equally similar to their mother in terms of gut microbiome composition.

### Mother-to-infant vertical transmission

Previous work on captive primates suggests that the effect of maternal parity on microbial function could be mediated by differences in vertical transmission between multi- and primiparous females, with primiparous females transferring more milk-oriented microbes to their offspring (via the milk) [10]. We tested if we could statistically detect evidence of vertical transmission between mother and offspring using fecal-fecal microbiome comparisons. We used a nonparametric resampling approach to test if mother-offspring pairs of fecal samples were more similar than expected by chance (i.e compared to when we match the immature sample with random adult female samples), as measured by the number of shared ASVs or beta dissimilarity. We predicted that vertical transmission would be strongest in early life (when infants are still nursing), thus we ran analyses using either (i) all samples (0-3 years, N=398 pairs) or focusing on (ii) young infants (<12 months of age, N=136 pairs) and (iii) old immatures (>18 months of age, N=201 pairs) separately. Using all pairs, we found that immatures shared 3.4% more ASVs (observed value=355, random value=343, *P*<1.0×10^-3^) and were 1.8% more similar compositionally (unweighted UniFrac dissimilarity: observed=0.55, random=0.56, *P*=1.0×10^-3^) to their own mother than with random adult females of the population (**Table S11**), potentially indicative of vertical transmission. However, unexpectedly, this signal was weaker and non-significant in the youngest infants (0-12 months: number of shared ASVs: observed=251, random=245, *P*=0.09 and unweighted UniFrac dissimilarity: observed=0.67, random=0.67, *P*=0.26; Figures 5C **and S12, Table S11**), and was strongest and significant in older juveniles (>18 months: number of shared ASVs: observed=412, random=398, *P*=2.0×10^-3^ and unweighted UniFrac dissimilarity: observed value=0.49, random value=0.50, *P*=1.0×10^-3^; Figures 5C **and S12, Table S11**). The finding of greater vertical transmission after, rather than before, nursing cessation suggests that these mother-offspring similarities were mostly mediated by non-nursing interactions and that milk vertical transmission may not be adequately captured by comparing infant and maternal fecal microbiomes.

Moreover, the ASVs shared between mother-infant pairs in the first 12 months of life were not the same ASVs found abundant in early-life (i.e., ASVs with a negative score on PC1) and therefore not related to nursing (Figure 5D**, Table S12**). For example, *Bacteroides fragilis* is found in 49% of infants <12 months but is only shared in 9% of mother-infant pairs. Instead, the most commonly shared ASVs among mother-infant pairs between 0-12 months tended to be ASVs characterizing later life (i.e., with positive scores on PC1), characteristic of older offspring and of adult females (Figure 5D**, Table S12**). Thus, mother-infant pairs share more bacteria and have more similar gut microbial community than expected by chance, but this shared community belongs to the typical adult microbiome of geladas, and is not specific to microbes functionally beneficial to processing milk during the early developmental period.

Since infants of primiparous females possessed more milk-oriented microbes (i.e., far from adult-like microbes), we also found that they shared fewer ASVs (ß=-74.5, *P*=0.01) and were more dissimilar to their mother (unweighted UniFrac: ß=0.07, *P*=0.03) in the first 12 months of life than infants born to multiparous females (Figure 5E**, Table S3**). However, this effect of greater dissimilarity in primiparous-infant dyads disappeared later in life (>18 months of age) when the effect of maternal parity was no longer detected (number of shared ASVs: ß=11.0, *P*=0.46, unweighted UniFrac: ß=-3.5×10^-3^, *P*=0.81) (Figure 5E**, Table S3**). This result shows that the effect of maternal parity on the offspring gut microbiome in the first 12 months of life is not mediated by stronger vertical transmission of milk-oriented microbes when using fecal-fecal comparisons.

## DISCUSSION

We provide a detailed description of the compositional assemblage and functional development of the infant gut microbiome in a nonhuman primate during the first three years of life. As expected, age was the strongest structuring factor of the diversity, composition, and function of the gut microbiome. Most microbial taxa had clear age-related trajectories and could be grouped into four main clusters that reflected progressive dietary transitions associated with weaning. In addition, our data show that maternal effects were an important factor modulating offspring gut microbiome both during nursing and after weaning.

The broad dynamic of microbial colonization in geladas presents many similarities with previous reports on humans [2,5,8] and other mammals ([10,37,73], but see[74]). We observed a low initial number of microbes and a rapid increase in microbial diversity in the first seven months of life, followed by more gradual changes in microbial composition until weaning (∼17 months). The fact that maximal microbial diversity was attained by the time infants reached 7 months, while the microbial community continued to evolve until weaning, suggests that numerous events of lineage extinction and *de novo* colonization continue to take place in the gelada gut until weaning. Similar to humans [5,42,93], it is the cessation of nursing rather than just the introduction of solid foods (which usually starts as early as the first few weeks after birth in geladas) that really drives the maturation of the developing gut microbiome to an adult-like composition. Indeed, weaning marks two important transitions that can have dramatic effects on the maturing gut microbiome. First, as milk is replaced by solid foods, the nutrient sources for host and microbes both change, altering the types of microbes that are likely to flourish. Second, weaning is accompanied by the loss of maternal-origin immunologic factors and milk-derived microbes [94], both of which can further alter the microbiome through processes of selective seeding. Shifts in gut microbiome composition and function closely followed progressive dietary transitions: gut bacteria that facilitate milk glycans and lactose utilization were dominant in the gelada microbiome during early infancy, while cellulolytic and fibrolytic bacteria that metabolize plant complex polysaccharides were dominant later in development as graminoids were progressively introduced in the diet [76]. Many of the early life colonizers were similar to those found in humans, such as *Bacteroides*, *Streptococcus*, *Faecalibacterium*, Lachnospiraceae, *Blautia*, *Clostridium*, *Veillonea*, *Escherichia-Shigella*, and Pasteurellaceae [48,82,95] which perhaps suggest a set of universal mammalian or primate infant microbial taxa. These early-life microbes work as a metabolic network that relies on cross-feeding between primary degraders (e.g., lactose-degraders such as *Streptococcus*) and secondary fermenters (e.g., lactate-utilizers such as *Veillonea*) to convert milk sugars into energy [96]. The functional enrichment in carbohydrate metabolism and fermentative pathways found in gelada infants is also typically observed in human newborns [2,5,55,97].

*Bacteroides*, and in particular *B. fragilis* [61,98], appear to be the primary microbial taxa involved in milk glycan degradation in geladas, as evidenced by their high abundance in early-life and the fact that they encode the enzymatic toolkits necessary to cleave complex milk glycans (e.g., fucosidase, sialidase, beta-galactosidase). These bacterial enzymes are critical for host nutrition, as mammalian hosts are unable to produce them and therefore cannot utilize milk glycans independently of gut bacteria [82]. In humans, this function is largely met by *Bifidobacterium*, a taxa commonly found in high abundance in breastfed humans that also breaks down milk glycans [8,48,99,100]; however this taxon was almost entirely absent in young geladas. In fact, variation in the dominance of *Bifidobacterium* and *Bacteroides* appears the norm at both the species and population level: several studies in mammals [10,37,72,101] and in some human populations [3,85,95,97,102] have noticed the absence of *Bifidobacterium* but abundance of *Bacteroides* in most or at least some nursing infants. *Bifidobacterium* and *Bacteroides* have different glycan-use profiles [61,62,97] linked to species and population differences in milk composition, particularly the structure and the relative abundance of different milk glycans [103–105].

The early-life microbiome of geladas was also characterized by a high number of potentially pathogenic bacteria known to cause enteric infection in human newborns and captive animals (*Clostridioides*, *Helicobacter*, *Clostridium*) [87–91] and several bacterial groups involved in the activation of the host immune system such as butyrate-producing (*Blautia*, *Faecalibacterium, Butyricicoccus, Butyricimonas*) and mucin-degrading bacteria (*Akkermansia*, *Ruminococcus gnavus* and *R. torques*). Collectively, this microbial profile suggests that immune function is a priority for gelada infants. Butyrate plays a key role in the maintenance of gut integrity [106,107] and protection against enteric infection [108]. This microbial metabolite is also an important immunoregulator via its action on intestinal macrophages [109,110]. Mucolytic bacteria play an essential role in mucus turnover [111] and contribute to an essential immune barrier protecting the underlying epithelium from luminal pathogens [111] and are thus strongly involved in immunity in early life. *Bacteroides* are also likely involved in regulation of intestinal immunity in early life [112,113]. *Bacteroides fragilis* in particular is directly involved in the maturation of the immune system by directing the production of regulatory T cells and ensuring a balance between Th1 and Th2 immunologic response [114–117]. Functional analyses revealed that the gut microbes were more strongly involved in host immunity during the nursing period, highlighting that microbial colonization plays an important role in priming of the host immune system in geladas.

We detected important compositional and functional signatures of microbial rearrangement around 10 months of age (i.e., 5-7 months before nursing cessation). *Bacteroides* decrease substantially in the gelada gut microbiome, while two other taxa, *Lactobacillus* and *Prevotella,* increase in abundance. *Lactobacillus* is a lactic acid bacterium that consumes lactose [118,119]. Its rise in abundance around the weaning transition indicates an increase in lactose availability in the colon, likely due to the loss of endogenous lactase of infants in the upper gut [120]. *Prevotella* is a keystone fiber-degrading bacterium typically enriched in individuals with a plant-based diet. In two other mammalian species (vervet monkeys: [10] and northern elephant seals (*Mirounga angustirostris*): [72]), *Prevotella* also increased in abundance during the weaning transition. The abundance of *Bacteroides* and *Prevotella* are generally inversely correlated in the gut, due to the trade-off between saccharolytic and proteolytic fermentation [121]. Thus, the growth of *Prevotella* closer to weaning might be related to the decrease in milk degrading bacteria (i.e., *Bacteroides*) and could be a good indicator of the transition from milk to solid food consumption in mammals [10]. These taxonomic changes were also accompanied by important functional changes in the metabolism of amino acids, vitamins, and cofactors, setting up the microbial activity characteristic of the adult gut.

Finally, our results highlight that early-life gut microbiome composition and functionality can be influenced by maternal effects, both during the nursing period, but also after weaning. During the first 12 months of life, we found that infants of primiparous mothers harbored more bacteria that were functionally relevant for processing milk sugars, which parallel recent findings in vervet monkeys [10]. The authors in that study hypothesized that infants of primiparous mothers may compensate for poor maternal investment by seeding more milk-oriented microbes that help infants extract more energy from milk [10]. In support of this, *B fragilis* was more abundant in the milk of low-parity vervet females, which resulted in higher abundance of milk-oriented microbes in the infant gut, which in turn promoted faster growth in low-parity infants [10]. In our study, vertical transmission – as assessed by fecal-fecal comparison of maternal and offspring communities – was not identified as the mechanism generating such a parity effect. First, we did not find evidence of vertical transmission in the first 12 months of life (infants and mothers did not share more ASVs than expected by chance during nursing). Second, the microbes that *were* shared by mother-offspring pairs were associated with processing grass rather than early life functions such as processing milk glycans or sugars. Third, infants from primiparous females actually shared fewer microbes with their mother than infants from multiparous females (since the detected shared microbes are later-life microbes). This result suggests that vertical transmission of early colonizers/milk-oriented microbes might be more strongly mediated by the direct transfer of milk microbiota in geladas [10,49,50,54] and, in contrast to reports in humans [42,44,55,56], not easily detected using fecal-fecal comparisons between infants and their mothers. In vervets, for instance, infants aged 2-5 days shared more bacterial strains with their mother’s milk than with their mother’s gut [10]. This parity effect could nonetheless result from host filtering processes coming from the offspring themselves [56]. Maternal microbiomes might be similar across parity status, but offspring of primiparous females might preferentially seed milk-oriented microbes from milk in response to poorer maternal energetic allocation. In the absence of milk samples, evidence for such mechanisms remains unclear in geladas.

Alternatively, the effect of maternal parity could reflect a faster pace of gut microbiome maturation for offspring born to multiparous mothers. The pattern of vertical transmission might be similar between primiparous and multiparous females, but offspring of multiparous females might share more microbes with their mother during the first 12 months of life because they are more mature for age (and because we only capture vertical transmission of grass-processing microbes). This interpretation is supported by the evidence that multiparous mothers wean their offspring about 5 months earlier than primiparous females in geladas (in our studied cohort and in absence of takeover: multiparous=17.1 months, primiparous=21.9 months). The greater similarity between multiparous mothers and their infants is thus more likely to be generated by accelerated gut microbiome development, suggesting that these infants are undergoing the weaning transition at a faster pace than their peers. Infants from multiparous females could be eating solid grass, gaining physical independence, and becoming socially integrated earlier than their peermates, all of which could explain greater microbial resemblance to mothers (and other adults). Behavioral and development data, such as infant growth, are needed to investigate this hypothesis of accelerated development and their consequences on offspring phenotype.

Somewhat surprisingly, we did find that immature gut microbiomes were more similar to maternal gut microbiomes than expected by chance after weaning regardless of the parity status of the mother. Such an effect has been previously documented in wild red squirrels (*Tamiasciurus hudsonicus*) [122] and chimpanzees (*Pan troglodytes*) ([74] but see [123]). Host genetics, or socially transmitted microbes, may facilitate maternal-offspring gut microbiome similarities beyond the early postnatal period [47]. A recent study in yellow baboons (*Papio cynocephalus*) found that the gut microbiome, including both abundant and rare taxa, is highly heritable [124] suggesting that the convergence of the gut microbiota between mother and offspring in geladas could be due shared genes. Alternatively, or additionally, the higher similarity in later life could be generated by high frequency of social contacts between mothers and offspring that extend past weaning. Primate mothers and offspring form preferential social bonds long after weaning, relationships that are characterized by a high degree of proximity, physical contact and grooming, and are likely to represent an enduring source of maternal microbial inoculation for offspring [58,125]. Further work is needed to understand the relative importance of these mechanisms in explaining mother-infant similarity during juvenility.

### Conclusion

Our results highlight that early-life gut microbiome composition and function can be influenced by maternal effects, both during nursing as well as after weaning. Maternal parity in particular was associated with the functional maturation of the microbiome in offspring, likely reflecting faster developmental pace of infants born to reproductively experienced mothers. As infants age and are weaned, they converge toward an adult-like gut microbiome that is more similar to the maternal gut microbiome than expected by chance. The long-term consequences of such microbially-mediated maternal effects remain unknown but could potentially influence phenotypic outcomes such as growth and immune function. Our work also emphasizes that early life vertical transmission, mediated in large part by milk transfer, may not be detected using fecal-fecal comparisons of maternal and infant communities and would ideally require data on the milk microbiome whenever possible.

## MATERIAL & METHODS

### Study population and study site

The data for this study were collected between Jan 2015 and Jan 2019 from a population of wild geladas living in the Simien Mountains National Park in northern Ethiopia (13°15′N, 38°00’E). Geladas live in multi-level societies, where several reproductive units (comprising a leader male, several adult females, their offspring, and occasionally 1–2 follower males) aggregate together during the day to forage and sleep together forming a “band”, sharing a homerange [80]. Since Jan 2006, the Simien Mountains Gelada Research Project (SMGRP) has collected behavioral, demographic, and genetic data on a near-daily basis from over 600 individuals living in 2 separate bands of the area. All gelada subjects were habituated to human observers on foot and were individually recognizable. Data were derived from 89 infants and juveniles aged between 0-3 years old and 83 adult females living in 23 different reproductive units. The date of birth of each infant was known within a few days’ accuracy. The reproductive state of each adult female was monitored daily and recorded as cycling (as indicated by the presence of sex skin swellings on the neck, chest, and perineum), lactating (if she had a nursing infant), or pregnant (the date of conception was inferred by removing 183 days from the date of birth of subsequent offspring) [81]. Records of female reproductive history were used to assign maternal parity status for each infant (first-time mother: primiparous or multi-time mother: multiparous) and to establish the date at which the mother resumed cycling following the infant’s birth, which we used to estimate the approximate age at weaning for each infant. For 8 infants, age at weaning began on the date of maternal death.

### Fecal sample collection

Fecal samples (N=525; 303 females, 222 male samples) from 89 immature geladas (i.e., infants and juveniles sampled pre-reproductive maturity; female: N=51; male: N=38, mean±SD=5.90±5.53 samples per individual, range=1-18) were collected opportunistically from 2015-2016, and then regularly from 2017 to 2018) during the development using targeted protocols (**Figure S1**). These samples come from individuals residing in 17 different reproductive units (mean±SD= 5.65±4.44 number of individuals sampled per unit, range=1-17). For a subset of immature samples (N=398 samples from 61 infants), we also collected a matched fecal sample from the mother (N=398 samples from 44 mothers) on the same day or on the following day of the infant sample collection. Fecal samples of known adult females in all reproductive states were also routinely collected (N=222 samples from 79 females) and were used to generate a random distribution of gut microbiome composition similarity between females of the population and immatures. Immediately upon defecation, approximately 1.5 g of feces was collected in 3 ml of RNA later [126] stored at room temperature for up to 2 months, and subsequently shipped to the University of Washington (UW). At UW, samples were stored at -80°C until the sequencing libraries were prepared.

### Maternal dominance ranks

Female dominance ranks were established using *ad libitum* and focal observations of agonistic interactions between all adult females belonging to the same unit with an Elo-rating procedure [127] implemented in the R package EloRating [128]. Agonistic interactions included physical aggression (hit, bite), chase, threats (vocal threats, non-vocal gestures), approach-avoid interactions (displacements) and submissive behaviors (fear bark, crouch, grimace). In geladas, agonistic interactions usually consist of a sequence of several behaviors in a row emitted and received by both parties. Since it can be difficult to establish the winner of each agonistic sequence, we consider each behavior of a sequence as a separate event and assign the winner and loser based on the directionality of the behavior. We obtained a daily Elo-score that we then averaged per month. Since Elo-scores can be sensitive to differences in sampling effort, we then converted this monthly Elo-rank into a monthly proportional rank and controlling for female group size (0=lowest-ranking females and 1= highest ranking female). In the analyses, we used maternal dominance rank during the month of the infant’s birth since we expect microbially-mediated maternal effects to be the strongest in the postnatal period (during nursing). However, we also investigated maternal rank during pregnancy and at the date of immature sample collection, which led to similar results (not reported here).

### Environmental data

The study area is located at 3200 m above sea level and is characterized as an Afroalpine grassland ecosystem, consisting of grassland plateaus, scrublands, and Ericaceous forests [129]. The climate in the Simien Mountains National Park exhibits marked inter- and intra-annual fluctuation in rainfall and temperature and can be broadly divided into 3 distinct seasons : a cold-dry season (Oct to Jan), a hot-dry season (Feb to May) and a cold-wet season (Jun to Sep) [130]. Fecal samples of immatures and adult females were collected across the year, with roughly equal coverage across seasons (406 in cold-dry, 426 in cold-wet and 313 in hot-dry season). Daily cumulative rainfall and minimum and maximum temperature are recorded on a near-daily basis by the SMGRP. Geladas are graminivorous, with up to 90% of their diet composed of graminoids [76]. They eat primarily graminoid leaves (i.e., grasses and sedges) all year long, but increase substantially their consumption of underground storage organs (rhizome, corms, roots) in the dry season, as above-ground graminoid leaves become less abundant [76]. A previous study established that the gut microbiome composition of adults shifts in response to environmental variation, in particular with cumulative rainfall which is a good proxy of diet. [79]. Thus, in all models we controlled the total cumulative rainfall over the 30 days prior to the date of fecal sample collection (as a proxy for grass availability) and the average minimum daily temperatures in the 30 days preceding the date of sample collection (as a proxy of thermoregulatory constraints).

#### 16S rRNA gene sequencing

We performed 16S rRNA gene amplicon sequencing on the immature and female fecal samples to establish gut microbial composition. We first extracted microbial DNA using Qiagen’s PowerLyzer PowerSoil DNA Isolation kit (Qiagen #12855) following standard protocols. We then amplified the hypervariable V4 region of the 16S rRNA gene using PCR primer set 515F and 806R from The Human Microbiome Project and a dual**-**indexing approach [131]. Details of the amplification protocol can be found in [79] (see also: https://smack-lab.com/protocols/). The libraries were then pooled in roughly equimolar amounts (each with their own unique indexing primer combination), spiked with 10% PhiX to increase library complexity, and sequenced together on a single Illumina NovaSeq 6000 SP 250 bp paired-end sequence flowcell at the Northwest Genomics Sequencing Core at the University of Washington, Seattle.

Data were processed using the Quantitative Insights Into Microbial Ecology 2 (QIIME2) platform [132] using the demux command to demultiplex raw reads and the DADA2 pipeline [133] to generate amplicon sequence variants (ASVs) feature tables. Forward and reverse reads were trimmed to 220 and 180 bases, respectively, to remove the low**-**quality portion of the sequences. Only samples with more than 20,000 reads were retained for analyses following observation of rarefaction curves. After filtering, trimming, merging, and chimera removal, we retained a total of 219,125,888 reads across the 525 immature fecal samples (417,382±645,328 reads per sample, range= 21,256- 7,976,983) and 293,003,271 reads across the 620 adult female fecal samples (472,586±869,181reads per sample, range= 20,068- 10,723,460). ASVs were taxonomically assigned using the q2**-**feature classifier in QIIME2 against version 132 of the SILVA database (updated December 2017) [134] based on 100% similarity.

#### Statistical analyses

The count and taxonomy files generated by QIIME2 were imported into R version 3.5.2 [135] using the qiime2R package [136]. We filtered the count table to retain only ASVs that had at least 500 reads total in the dataset to eliminate potentially artifactual sequences. With this filtering criteria, only 3,884 ASVs remained (out of the 29,686 initially observed). In total, 3,784 different ASVs were found across the 525 immature fecal samples (mean±SD number of ASVs per sample: 728±261, range: 65-1498), while the 620 female samples contained 3,679 ASVs (mean±SD number of ASVs per sample: 829±248, range: 98-1761). Most ASVs could be taxonomically assigned to the phylum (100%), class (99%), and order levels (99%), with assignments decreasing substantially at the family (88%) and genus (63%) levels.

##### Alpha-diversity analyses

We calculated three complementary metrics of alpha diversity for each sample: the observed richness (the total number of unique ASVs per sample), Shannon Index (taking into account both richness and evenness in abundance of ASVs), and Faith’s phylogenetic diversity (a measure of the diversity of phylogenetic lineages within a sample) using the “phyloseq” [137] and “picante” package [138]. To assess which predictors affected immatures’ gut microbial alpha diversity, we used generalized additive mixed models (GAMMs) with the ‘mgcv’ package in R [139]. Such models allow fitting of a nonlinear relationship between the response variable and the fixed effect (by adding a smooth term), such as between alpha diversity and immature age (Figure 1C). Fitted predictors included: immature age at the date of fecal sample collection (modeled as a smooth term), immature sex, the parity status of mother, maternal dominance rank in the month of infant’s birth, cumulative monthly rainfall, average monthly minimum temperature and the log-transformed sequencing depth (i.e., the number of reads per sample). The use of rarefaction (i.e., subsampling of the read counts in each sample to a common sequencing depth) has been strongly discouraged on microbiome dataset because it discards too much sequencing information and leads to high rate of false positives [140], so we calculated alpha diversity on raw counts but controlled for sequencing depth in our model. Graphical representation of alpha diversity metrics are nonetheless displayed using a rarefied dataset at 20,000 reads. Individual identity and unit membership were included as random effects. Model residual checks were performed using the qq.gamViz and check.gamViz functions. Given that GAMMs models can not accommodate the test of the interaction between a smooth and fixed term, we ran those models including all immature samples or on only young infants (0-12 months) to test for the significance of maternal effects in early life (i.e., when the infant is still nursing).

To quantitatively assess the age at which alpha diversity reaches a plateau (i.e., converges to adult-like pattern), we used quadratic plateau models (formula: y ∼ (a + b * x + c * I(x^2)) * (x <= -0.5 * b/c) + (a + I(-b^2/(4 * c))) * (x > -0.5 * b/c)) fitted using the nlsfit() function of the easynls package [141] and extracted the critical point of inflexion and r-squared of the optimized model (i.e., with the values of a, b and c fit best the data). Since it is not possible to control for covariates in those analyses (e.g., sequencing depth), we ran those models on a rarefied dataset at 20,000 reads.

##### Beta-diversity analyses

Beta-diversity (between-sample dissimilarity in composition) was computed as the Aitchison distance [142], which is simply the Euclidean distance between samples after centered log-ratio (clr) transformation of the raw counts (a pseudo-count was added to the zeros using the imputation based on a Bayesian-multiplicative replacement from the cmultRepl() function in the package zCompositions [143]). The clr transformation allows us to account for differences in sequencing depth between samples and is a better practice than rarefaction of the counts [144]. Principal components analysis (PCA) on the Aitchison dissimilarity matrix (function “prcomp”) was used to examine how immatures samples clustered by age. We extracted the loading scores for each ASV onto the first Principal component (PC1) of the PCA to determine which specific ASVs have the highest influence on the clustering by age of samples. A quadratic plateau model was implemented to find the age at which Aitchison beta diversity reaches a plateau.

Permutational Multivariate Analysis of Variance (PERMANOVA) was then carried out on the Aitchison dissimilarity matrix using the adonis2 function in the vegan package [145] (with 10,000 permutations) to test for associations among gut microbial beta-diversity and the variables of interest (immature age, sex, maternal parity, maternal rank, environmental variables, the log-transformed sequencing depth, and unit membership). Individual identity was included as a blocking factor (“strata”) to control for repeated sampling among individuals. PERMANOVA models were run when including all immatures samples or on only young infants (0-12 months) to test for the significance of maternal effects in early life. We also replicated those PERMANOVA analyses using more classical measures of beta diversity (unweighted and weighted UniFrac dissimilarity) on a rarefied dataset at 20,000 reads and found essentially similar results (**Table S13**).

##### Mother-infant comparison of gut microbiome composition

To assess similarity in gut microbiome composition between mother and offspring, we calculated (1) the number of shared ASVs across maternal and immature communities, and (2) the beta diversity dissimilarity (unweighted and weighted UniFrac distances) between the matched infant-mother fecal samples collected the same day (N=398). The dataset of immature and mother fecal samples was rarefied at 20,000 reads to calculate these metrics since sequencing depth is likely to affect the similarity between paired samples. Quadratic plateau models were implemented on the three metrics to identify the age at which infants converged toward the maternal (i.e., adult-like) gut microbial composition. To assess which predictors affected the compositional similarity between mother-offspring pairs, we used GAMMs to model those three metrics as a function of immature age (as a smooth term), immature sex, maternal parity and maternal dominance rank, climatic variables (cumulative monthly rainfall and average monthly minimum temperature), while individual identity and unit membership were included as random effects. These GAMMs were also run separately on young infant samples (<12 months) or only on old immatures (>18 months) to assess how the strength of vertical transmission varies with maternal traits.

##### Individuality of the microbiomes in immatures

To capture the compositional divergence between immature samples, we calculated a measure of “individuality” of the microbiomes among the 525 immature samples, as defined in [146], which corresponds to the beta diversity dissimilarity value between a sample and the most similar sample (i.e., the minimum pairwise values from a beta diversity dissimilarity matrix, based on unweighted and weighted UniFrac metrics). The higher the value, the more distinct the gut microbiome composition is from all other immature samples in the cohort. This was calculated using the rarefied dataset at 20,000 reads.

##### Age-associated changes in microbial taxonomic composition

To identify the microbial taxa that vary significantly in abundance as immatures age, we used a statistical framework that is commonly used to analyze time series (and, in our case, longitudinal dataset). Autoregressive Integrated Moving Average (ARIMA) models allowed us to model and test for chronological trends in temporal data [147]. First, raw microbial counts were aggregated at the family or genus level, normalized using a clr-transformation, and z-transformed per taxon (i.e., across samples) to correct for variation in library size and unaccounted variance due to other covariates. Only microbial families or genera ≥ 0.01% relative abundance across the samples were selected for further analyses. Second, the counts were averaged across samples belonging to the same chronological age and converted into z-ordered objects (using R package zoo [148]) and into time series objects. Formatted time series were then analyzed using auto.arima (from the forecast R package [149]), using stepwise search and Akaike Information Criterion (AIC) to select the best model. This algorithm scans a wide range of possible ARIMA models and selects the one with the smallest AIC. ARIMA models that exhibited significant non-stationary trends (as opposed to unstructured “noise” fluctuations indistinguishable from stationary data) were selected following the criteria in [147]: (1) the difference order from stationary was higher than zero, and (2) at least one autoregressive (AR) and moving average (MA) coefficient was included in the model. LOESS regressions were then fitted to re-predict the count of each taxon as a function of age.

We then grouped bacterial taxa into clusters based on similarities in age-associated abundance trajectories. Pairwise distances between microbial taxa trajectories (i.e., the predicted values of the LOESS regression) were computed using correlation coefficients as a distance measure [150], and hierarchical clustering was performed using the complete method (using the function hclust from the stats R package). The optimal number of clusters was determined using the Elbow method (i.e., choosing a number of clusters so that adding another cluster does not highly improve the total within-cluster sum of squares) [151]. Results of hierarchical clustering were visualized using the R package heatmap3 [152] to provide an overview of gut microbiome composition changes with age.

##### Age-associated changes in microbial functional composition

To predict the microbial functional metagenomes of each sample from 16S rRNA data, we used Phylogenetic Investigation of Communities by Reconstruction of Unobserved States 2 (PICRUSt2) v.2.1.3-b software [84] with default options (picrust2_pipeline.py). We then computed the relative abundance of Kyoto Encyclopedia of Genes and Genomes (KEGG) Orthologs (KOs) (agglomerated at level 2 or 3 of the BRITE map) and of Enzyme Commission (EC) numbers for each sample. The accuracy of the PICRUSt2 predictions for each sample were assessed by calculating the weighted Nearest Sequence Taxon Index (NSTI) score, a measure of how similar the bacteria from the sample are to reference genome sequences. The mean±SD NSTI score across the 525 immature samples was 0.49 ± 0.19 (range: 0.01-0.89).

The age-related temporal trajectory of each KO pathway and EC was assessed using ARIMA models in a similar fashion than described above. The only difference is that the raw metagenome counts were transformed into relative abundance (instead of clr transformed). Only microbial pathways ≥ 0.01% relative abundance across the samples were included. Hierarchical clustering was used to group the pathways with similar aging trajectories.

##### Maternal effects on offspring’s gut microbiome development

We examined how maternal traits (dominance rank, parity) were associated with differences in offspring gut microbiome (1) composition (at the family and genus levels) and (2) function (KO pathways at level 2 or 3 and EC numbers) using GAMMs models. We modelled the relative abundance of each taxon and each functional pathway as a function of maternal parity and maternal dominance rank in the month of infant’s birth, while controlling for immature age (as a smooth term), immature sex, climatic variables (cumulative monthly rainfall and average monthly minimum temperature. For (1), the logarithm of the relative abundance of each taxon was fit (adding a pseudo-count of 0.001% to include zero counts). In all models, individual identity and unit membership were included as random effects. P-values were adjusted for multiple hypothesis testing by calculating the Benjamini-Hochberg FDR multiple-test correction. Only taxa that had an average relative abundance across samples ≥ 0.01% were tested. Given the number of metabolic pathways and the correction of p-values for multiple testings, only pathways that had an average relative abundance across samples ≥ 0.10% were tested. Taxa or functional pathways with a p-value < 0.05 were considered statistically significant. These analyses were run including all immatures samples (0-3 years), only young infant samples (<12months) or only old samples (>18 months).

##### Mother-to-infant vertical transmission

To assess if maternal and infant gut microbiome communities were more similar than expected by chance, we took a resampling approach (with 1000 repetitions) to compare the number of shared ASVs and beta diversity dissimilarity metrics (unweighted and weighted UniFrac) between (1) actual mother-infant matched samples (the observed value) and (2) random pairs of fecal samples of an infant and an adult female of the population (the random distribution). Since mother-infant pairs always shared the same social unit and were always collected 0-1 day apart (i.e., in the same season), we needed to match the random female samples accordingly to avoid introducing consistent bias in the random distribution. The random matching was thus done by either matching the infant sample to (i) a female of the same unit (to control for higher similarity only due to sharing the same social group) or (ii) a female sample collected in the same season (to control for higher similarity only due to seasonality). We did not have enough female samples to match by both criteria simultaneously. After we created the set of random pairs, we use GAMMs to compare the observed and random distribution of the metrics (number of shared ASVs or beta diversity dissimilarity) (response variable) by fitting a variable (“type of pairs?”) coding whether the value comes from an actual mother-offspring pair (1=observed) or a random infant-female pair (0=random), and controlling for immature age (as a smooth term) and immature sex. Infant and female identity were included as random effects to account for repeated observations of the same individuals. We extracted the estimate of the “type of pairs?” variable for the model and re-ran the model on a different set of random pairs (1000 times in total). We thus obtained a distribution of 1000 estimates for the “type of pairs?” variable. We report the exact p-value (calculated as the proportion of models with positive estimates for the number of shared ASVs and the proportion of models with negative estimates for beta dissimilarity) and the 95% confidence interval of the estimates of the “type of pairs?” variable. Fecal samples were rarefied at 20,000 reads to control for differences in sequence depth between infant and female samples. These analyses were run including all immatures samples (0-3 years), only young infant samples (0-12 months), or only old immatures (>18 months) to compare the strength of the effect among the different age categories.

To examine the nature of the shared microbes between mother and offspring in early life (when infants are <12 months), we extracted all ASVs in common between the 136 mother-offspring pairs (on the rarefied dataset). For each ASV found in the young infant samples (<12 months, N=3,402 ASVs total), we simply computed its relative abundance and prevalence across samples and how many pairs shared this given ASV. We then plotted the loading score of the ASV on PC1 of the beta diversity ordination (PC1 correlates strongly with age, so ASVs with the most negative versus positive loading scores are found in early versus later life) according to the percentage of mother-offspring pairs sharing this ASV.

## ACKNOWLEDGEMENTS

We thank the Ethiopian Wildlife Conservation Authority (EWCA), along with the wardens and staff of the Simien Mountains National Park for permission to conduct research and ongoing support to our long-term research project. We are also very grateful to the Simien Mountains Gelada Research Project field team for their help with field data collection, particularly our primary data collectors (Esheti Jejaw, Ambaye Fanta, Setey Girmay, Atirsaw Adugna, Dereje Bewket) and research assistants in the field (in particular Liz Babbitt, Maddie Melton). We would like to thank Marina Watowich and Dr. Kenneth Chiou for helpful discussions on ARIMA analyses.

## AUTHORS’ CONTRIBUTIONS

Conceptualization, Data curation, Formal analysis, Investigation, Visualization: A.B, A.L., N.S.M; Methodology/Laboratory work: A.B., S.S., A.M., Writing – Original Draft: A.B, A.L., N.S.M; Writing – Review & Editing: all authors; Funding Acquisition: A.L., N.S.M., J.C.B., T.J.B., L.R.; Supervision: A.L. and N.S.M.

## AVAILABILITY OF DATA AND MATERIALS

All 16S sequence data used in this study are available at the NCBI Sequence Read Archive under BioProject ID: PRJNA772269 (http://www.ncbi.nlm.nih.gov/bioproject/772269) for the immature samples **[temporary note: the sequences coming from these samples have been deposited on NCBI, but will only be released upon acceptance of the manuscript]** and PRJNA639843 (https://www.ncbi.nlm.nih.gov/bioproject/PRJNA639843) for the adult female samples. Data (including the ASV table and metadata) and R code to reproduce all the analyses are available at: https://github.com/GeladaResearchProject/Baniel-et-al_2022_Infant-gut-microbiome.

