## Supplementary information (text, tables, figures) for "Maternal effects on early-life gut microbiome maturation in a wild nonhuman primate": Supporting Information.docx

**This PDF file includes:**

Supplemental Results (1 and 2)

Supplemental Figures (S1 to S10)

Legends of supplemental Tables (S1 to S13)

**Supplemental Results**

**Supplemental results 1. Inconsistencies between age-related abundance trajectories of taxa.** Several microbial families and genera appeared to be classified in an irrelevant cluster (as based on clr-transformed counts) when comparing to raw abundance data (i.e. relative abundance, here on a log scale). At the family level, this was the case for Ruminococcaceae (Figure below, panel A) and Eggerthellaceae (Figure below, panel B) that are classified in the early-life cluster (i.e. decrease in abundance with age), despite showing a clear trend to increase with age. At the genus level, this was the case for *Intestinibacter* (Figure below, panel C) and *Ruminiclostridium 5* (Figure, D) that are classified in the early-life cluster (i.e. decrease in abundance with age), despite showing no real age-associated trajectory. Those misclassifications seem to happen especially when there is a lot of variability in the abundance of the taxon in early life. For example, Ruminococcaceae is mostly absent in early life but increases very rapidly with age (the adult-like abundance is reached at 5 months) and a couple of young infants (1-2 months) harbor already a particularly high abundance of it. These samples become outliers in the clr-transformed counts distribution and seem to drive the classification in the decreasing trajectory.

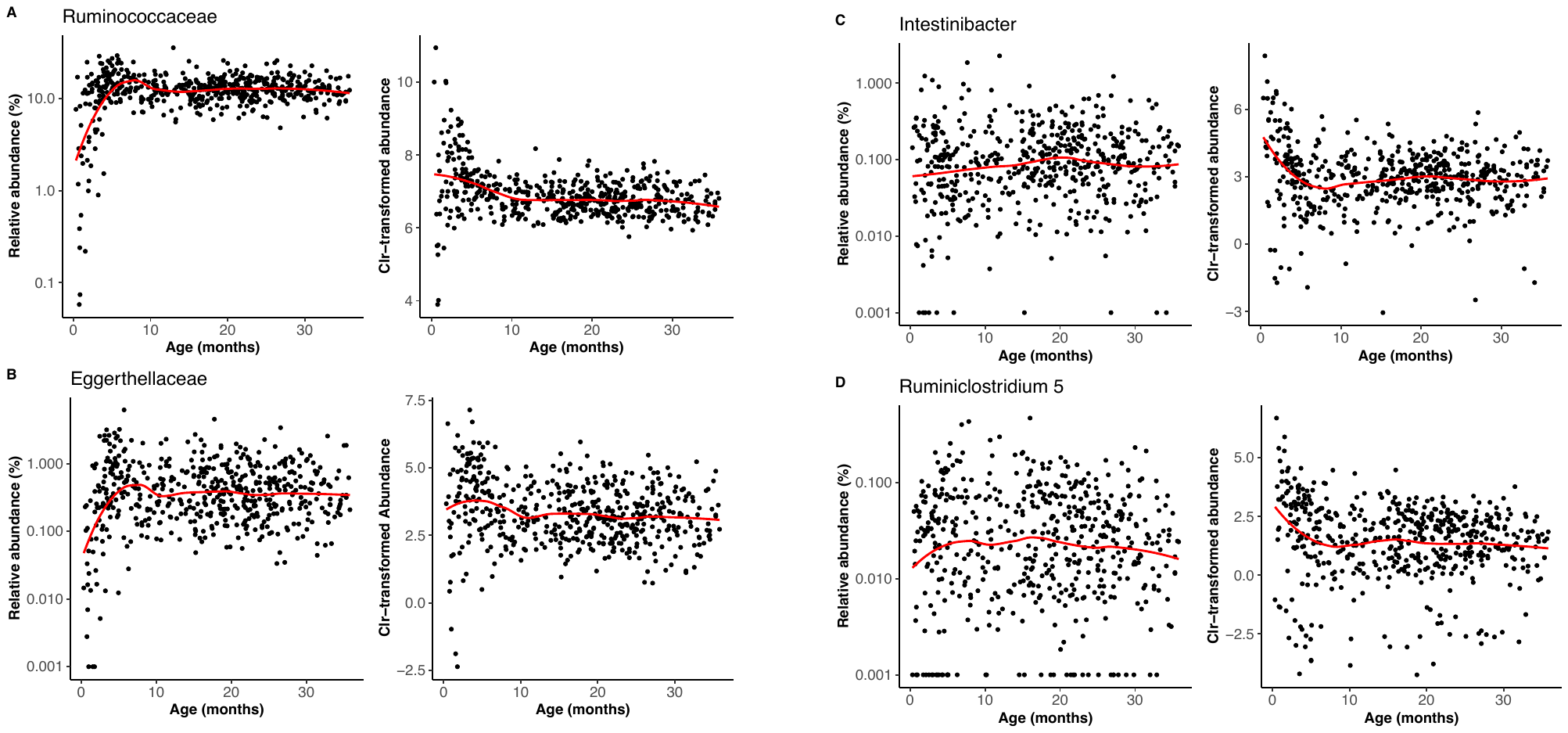

**Supplemental results 2. Effect of maternal parity on offspring’s gut microbiota composition.**

We could detect an effect of maternal parity on offspring’s gut microbiota at the functional level, but not at the taxonomic level. In fact, several taxa were differentially abundant according to maternal parity at the family and genus levels (or displayed a non-significant trend) before FDR correction of the p-values. Upon examining those taxa, we can notice that many of them are involved in milk digestion (**Figure S11)**. At the family level, infant born to primiparous females tend to have more Lachnospiraceae (β=0.15, p-value=0.011), Bacteroidaceae (β=0.44, p-value=0.076) and Clostridiaceae 1 (β=0.24, p-value=0.086) before FDR correction of p-values. At the genus levels several genera from the Lachnospiraceae family (*Roseburia*, *Lachnoclostridium*, *[Ruminococcus] torques group*, *Anaerostipes*) were significant significant (p<0.05 before FDR correction), as well as other genera characteristic of early life (*Alistipes*, *Erysipelatoclostridium*, *Bacteroides*). *[Ruminococcus] torques group* notably contains the *R. lactaris* species that is one of the most characteristic early life ASV om geladas and that is involved in lactose fermentation [[1,2]](https://paperpile.com/c/An11lE/KWYY+TeE3). *Bacteroides* contains *B. fragilis* that metabolize milk oligosaccharides [[3–5]](https://paperpile.com/c/An11lE/pe1y9+ACGjP+4oGwy). We thus conclude that milk-degraders exert small additive effects on the gut of infants of first-time mothers that are only detected at the functional level (which consider functional redundancy between taxa).

**Supplemental Figures**

**Figure S1.** Sampling design of the 525 fecal samples collected from 89 different immatures (y-axis) over the first 3 years of life (x-axis), together with information regarding whether the individual is considered weaned or not at the time of sample collection (using maternal cycling resumption as a proxy of nursing cessation).

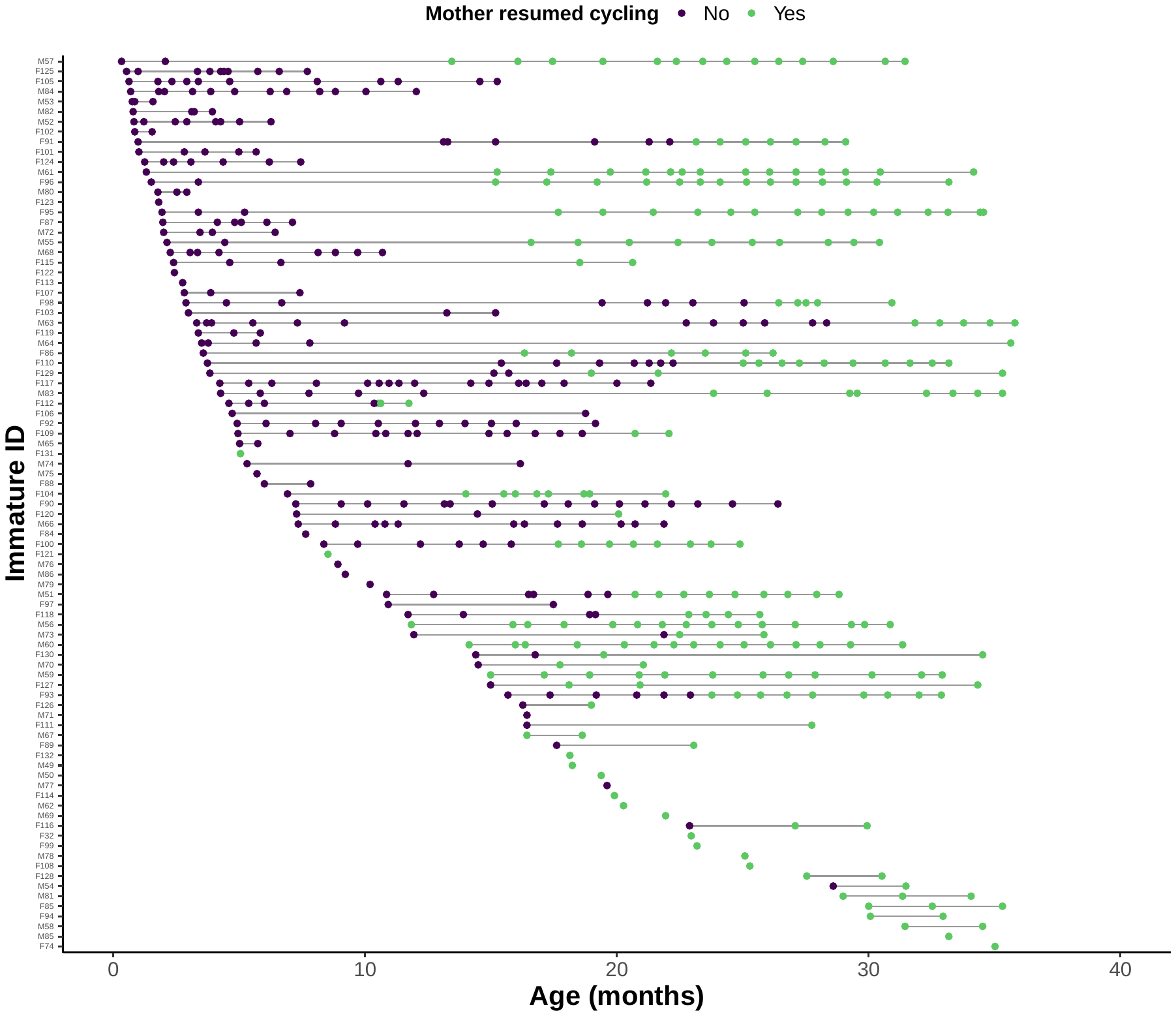

**Figure S2.** Age-associated pattern of alpha diversity within samples, as measured by (A) the observed richness (i.e. the number of Amplicon Sequencing Variants, ASVs, in a sample) and (B) the Faith’s phylogenetic diversity (PD) index. Vertical lines represent the critical points of inflexion (calculated using quadratic plateau models) representing the age at which alpha diversity converges to an adult-like value for the respective metrics. The dataset was rarefied at 20,000 reads for the figure.

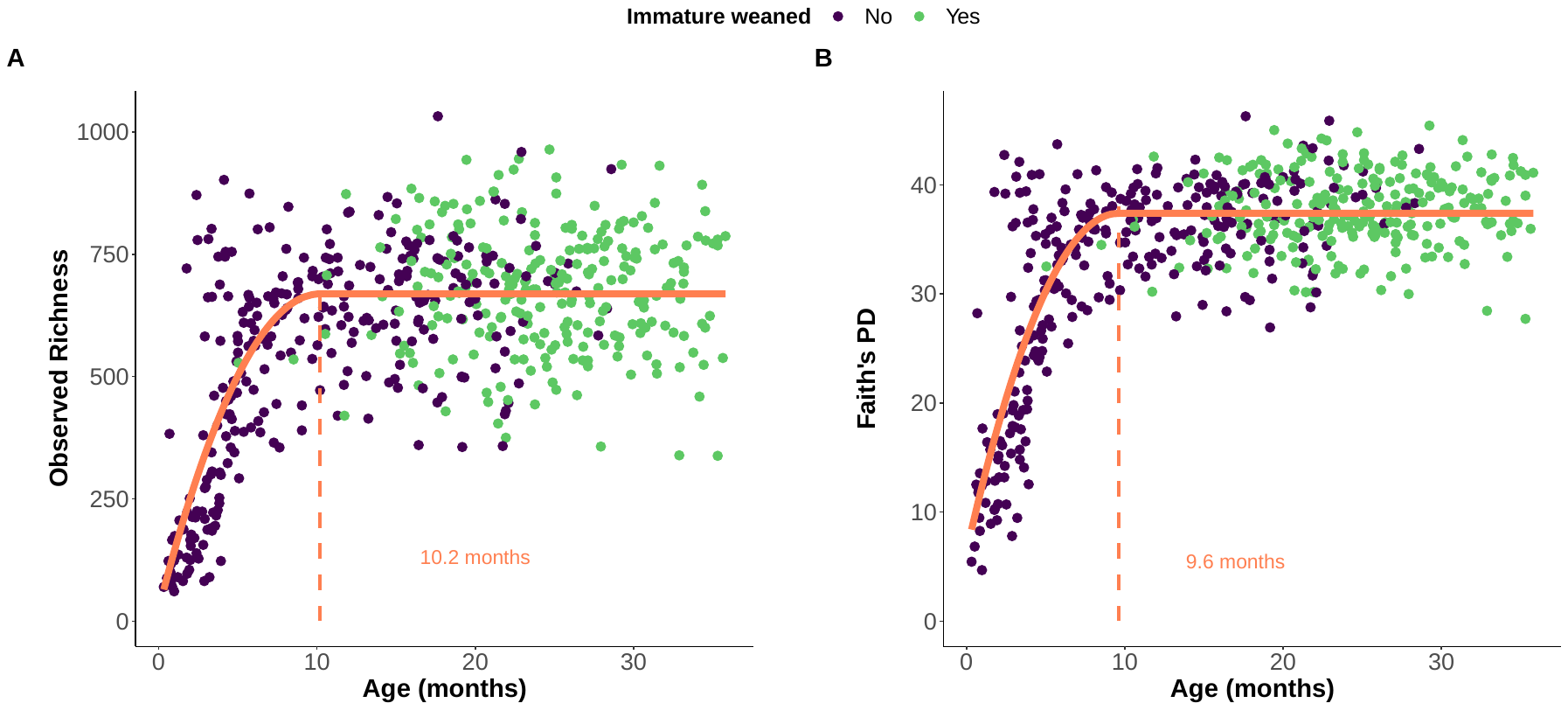

**Figure S3. Compositional maturation of the immature gut microbiome relative to the maternal gut microbiome.** Comparison of gut microbiome composition between mother and offspring, as assessed using 398 matched infant-mother fecal samples collected the same day. Two different metrics of beta diversity (unweighted and weighted UniFrac dissimilarity) – capturing broad difference in overall composition between the 2 samples – were calculated. The vertical lines represent the critical points of inflexion (calculated using quadratic plateau models) identifying the age at which the immature microbiomes converge to a community that is close from mothers. The dataset was rarefied at 20,000 reads for the calculation.

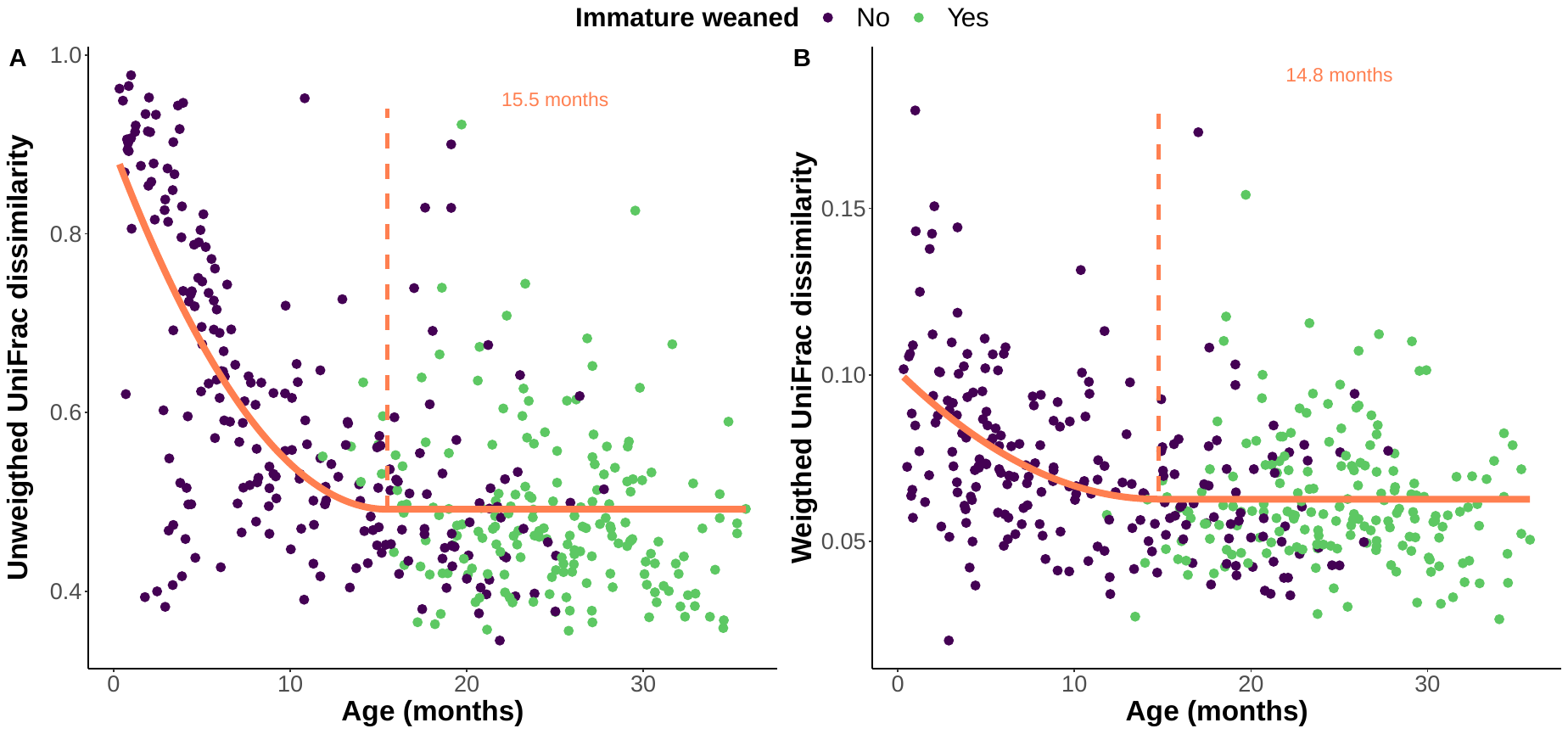

**Figure S4. Age-associated changes in microbial composition at the genus level.** Heatmap of the microbial genera exhibiting a significant chronological trend as a function of age (fitted values from ARIMA models and predicted using LOESS regression per taxa as a function of age). Values represent z-score normalized counts after centered-log-ratio (clr) transformation. Hierarchical clustering was used to group these age-dependent trajectories into four clusters exhibiting similar chronological trends. Color bars on the left side represent the delimitation of the clusters. Taxa (i.e. rows) are ordinated on the heatmap using correlation as distance function. All microbial genera above 0.01% abundance were analyzed (N=153) and 140 displayed a significant trend. On the right, the names of the genera above 0.1% abundance are displayed (per cluster).

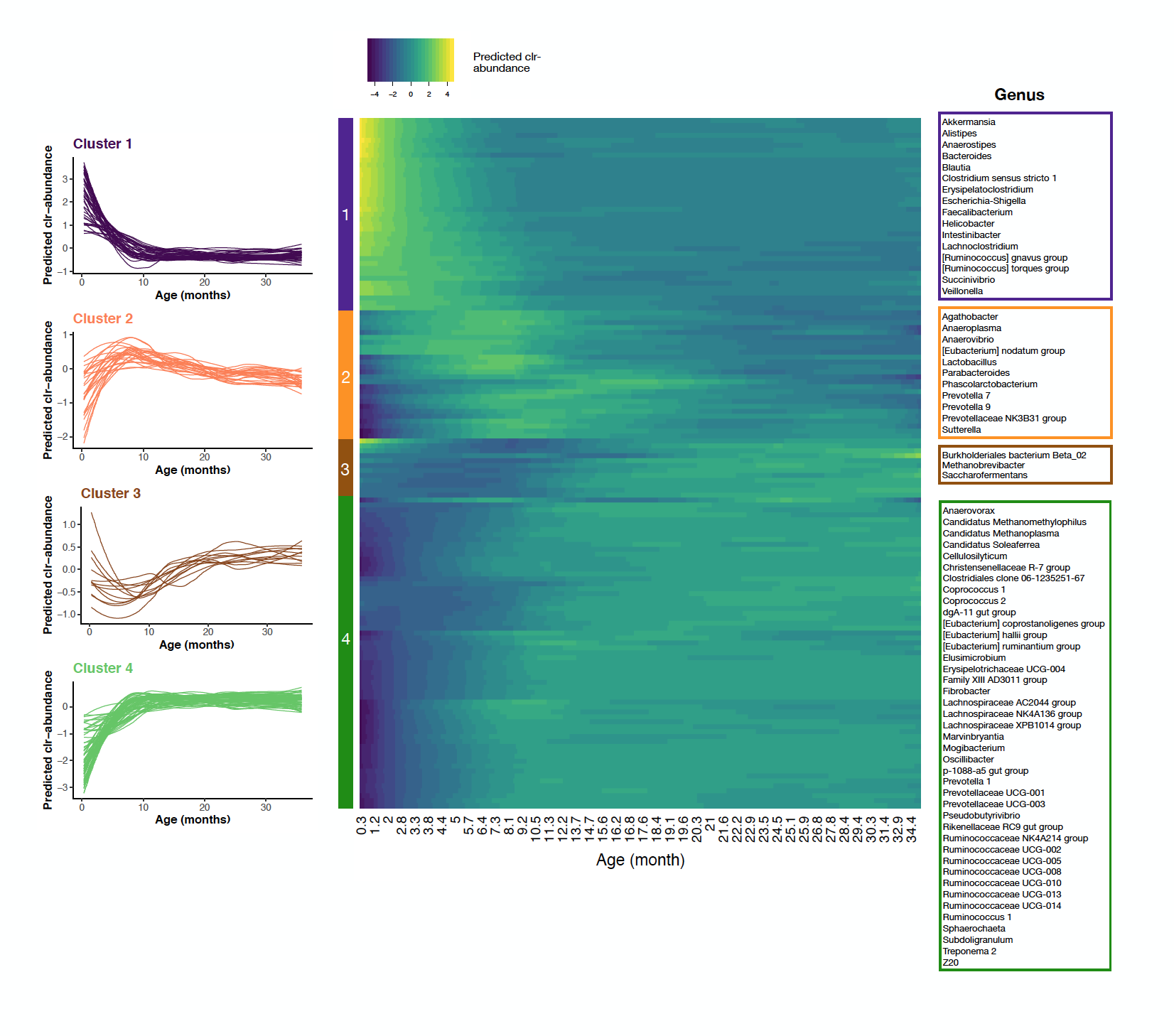

**Figure S5.** **Age-associated changes in the functional profile of the gut microbiome at level 2 of the KEGG orthologs (KO).** Heatmap of the predicted KO pathways exhibiting a significant chronological trend as a function of age (fitted values from ARIMA models and predicted using LOESS regression per pathway as a function of age). Values represent z-score normalized counts after relative abundance transformation. Hierarchical clustering was used to group these age-dependent trajectories into four clusters exhibiting similar chronological trends. Color bars on the left side represent the delimitation of the clusters. Pathways (i.e. rows) are ordinated on the heatmap using correlation as distance function. All pathways above 0.01% abundance were analyzed (N=37) and 33 displayed a significant trend and are represented.

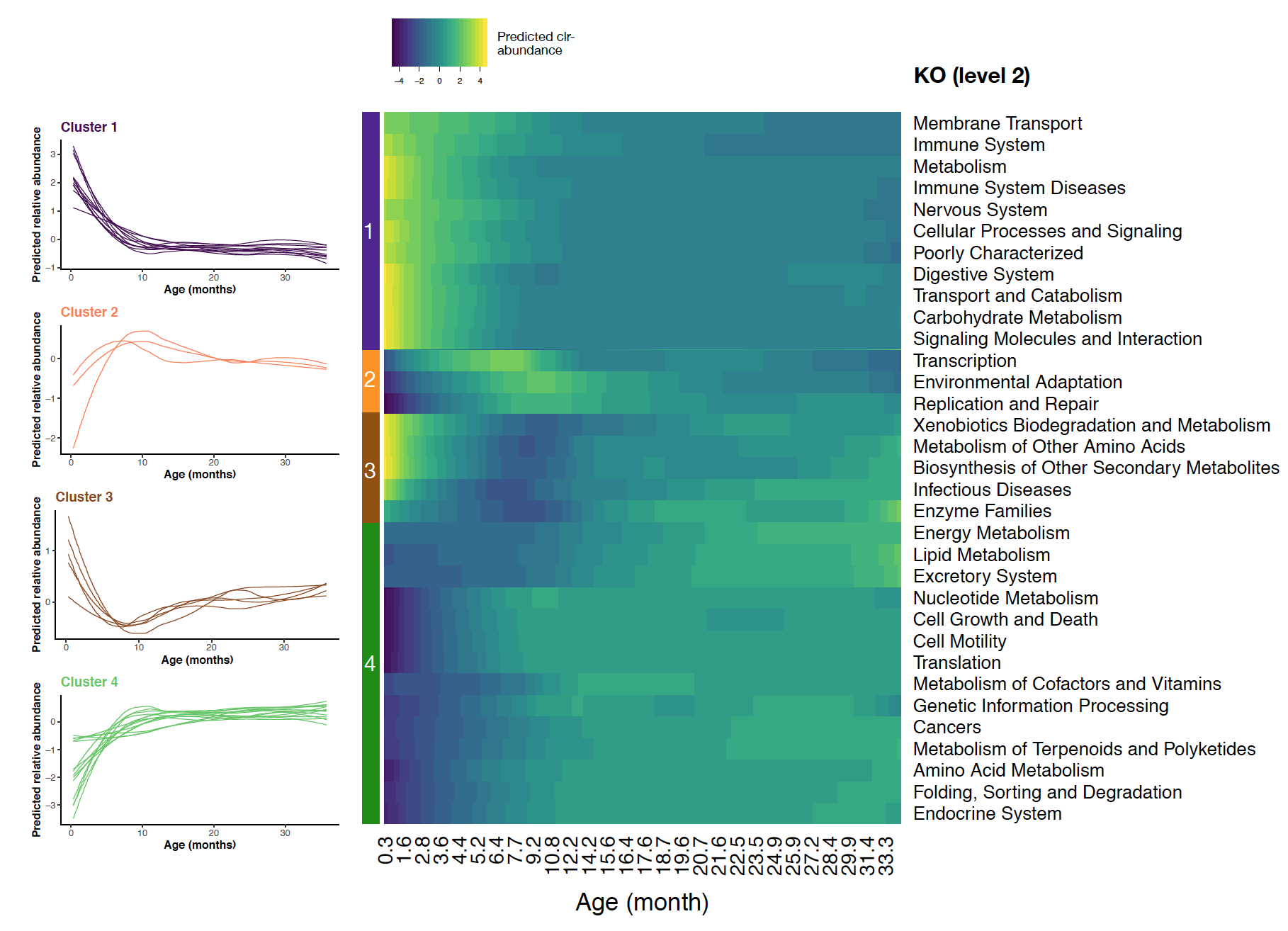

**Figure S6. Age-associated changes in the functional profile of the gut microbiome at level 3 of the KEGG orthologs (KO).** Heatmap of the predicted KO pathways exhibiting a significant chronological trend as a function of age (fitted values from ARIMA models and predicted using LOESS regression per pathway as a function of age). Values represent z-score normalized counts after relative abundance transformation. Hierarchical clustering was used to group these age-dependent trajectories into four clusters exhibiting similar chronological trends. Color bars on the left side represent the delimitation of the clusters. Pathways (i.e. rows) are ordinated on the heatmap using correlation as distance function. All pathways above 0.01% abundance were analyzed (N=208) and 189 displayed a significant trend. The heatmap only shows the 75 significant pathways above 0.4% relative abundance.

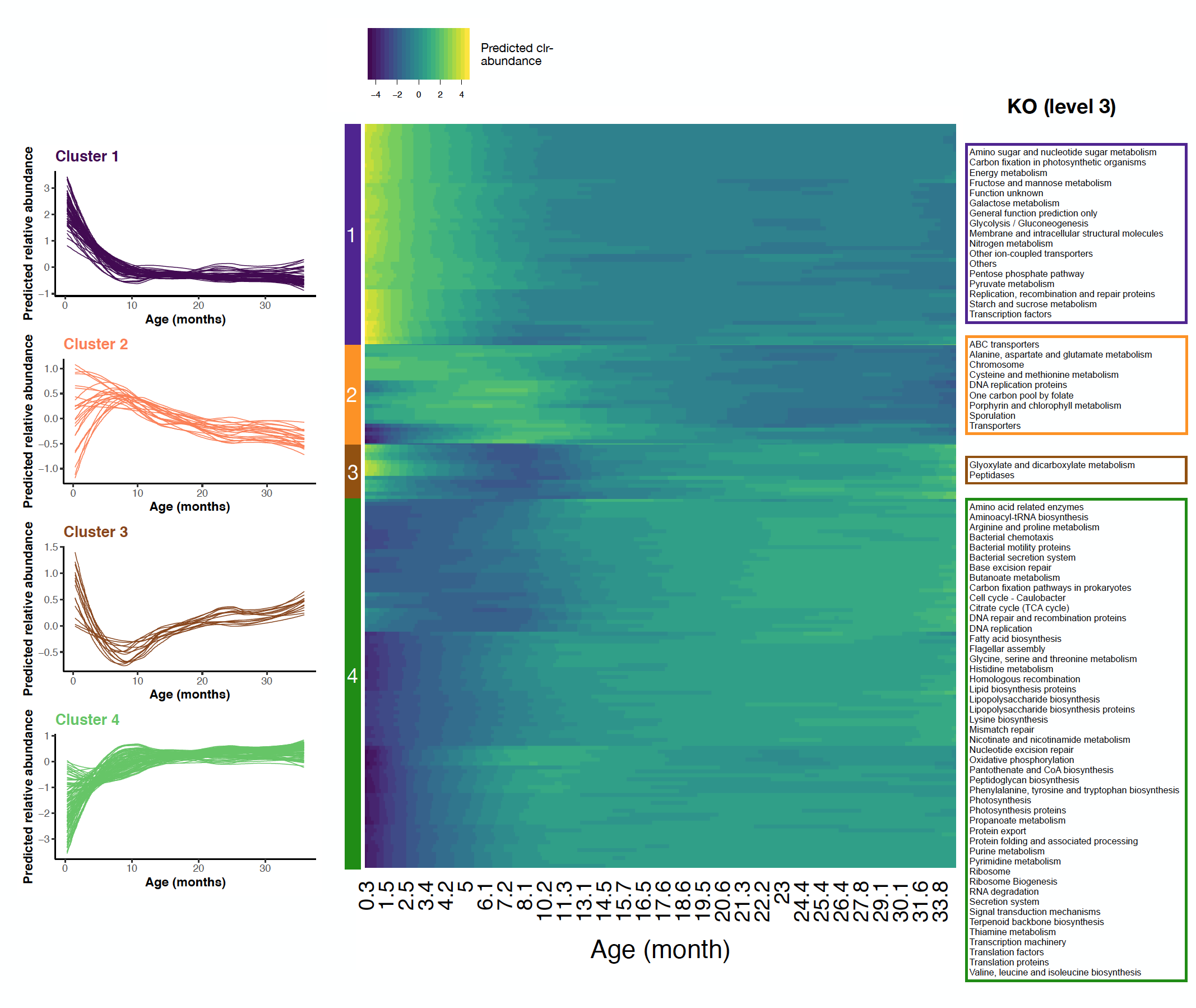

**Figure S7. Age-associated changes in the functional profile of the gut microbiome in Enzyme Commission (EC) numbers.** Heatmap of the predicted EC numbers exhibiting a significant chronological trend as a function of age (fitted values from ARIMA models and predicted using LOESS regression per pathway as a function of age). Values represent z-score normalized counts after relative abundance transformation. Hierarchical clustering was used to group these age-dependent trajectories into four clusters exhibiting similar chronological trends. Color bars on the left side represent the delimitation of the clusters. EC (i.e. rows) are ordinated on the heatmap using correlation as distance function. All EC above 0.01% abundance were analyzed (N=869) and 812 displayed a significant trend. On the right, the names of the EC above 0.17% abundance are displayed (per cluster). In cluster 1, the EC with a “*” had less than 0.17% abundance, but are highlighted in the text as functionally important to break down milk glycans moieties.

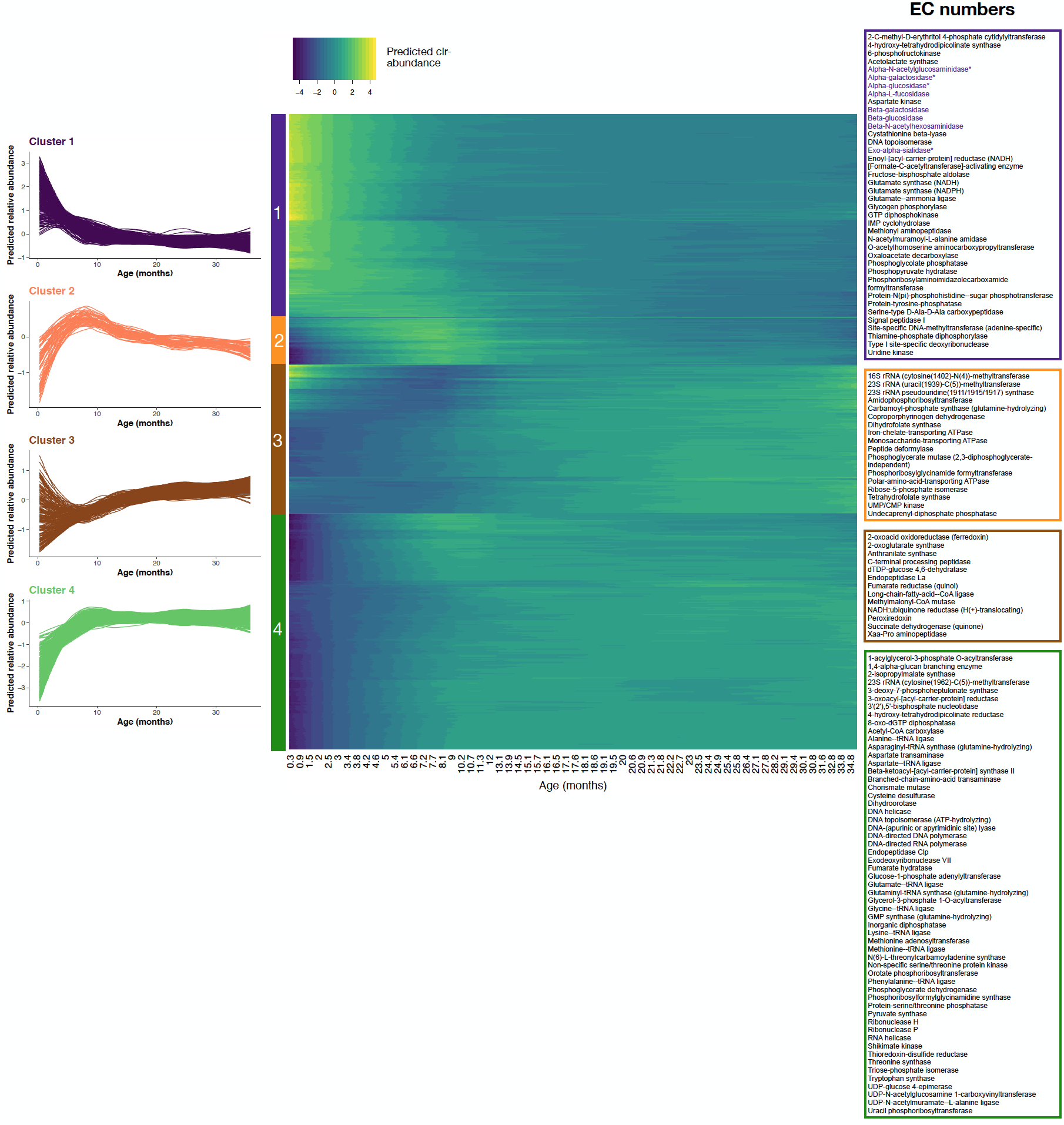

**Figure S8. The early-life gut microbiome encodes key enzymes able to break down milk glycans.** Milk glycans are complex oligosaccharides formed by a lactose core that can be elongated by a combination of moieties, such as fucose, sialic acid (N-acetyl-neuraminic acid), or N-acetylglucosamine. The enzymatic repertoire of the gut microbiota in early life was enriched in the four key glycoside hydrolases that decompose lactose into glucose (alpha-glucosidase and beta-glucosidase) and galactose (alpha-galactosidase, beta-galactosidase also called lactase), as well as in enzymes able to remove fucose (alpha-L-fucosidase), sialic acid (exo-alpha-sialidase) and N-acetylglucosamine (alpha-N-acetylglucosaminidase, beta-N-acetylhexosaminidase) moieties. The figure shows the relative abundance of 6 of these important enzymes, as a function of age (the averaged trajectory for each EC is obtained by fitting a LOESS regression across all samples of the raw relative abundance datapoints) and the distribution of the metagenomic contribution of the bacteria to each enzyme at the genus level. The averaged trajectory for each EC was obtained by fitting a LOESS regression across all samples of the raw relative abundance datapoints.

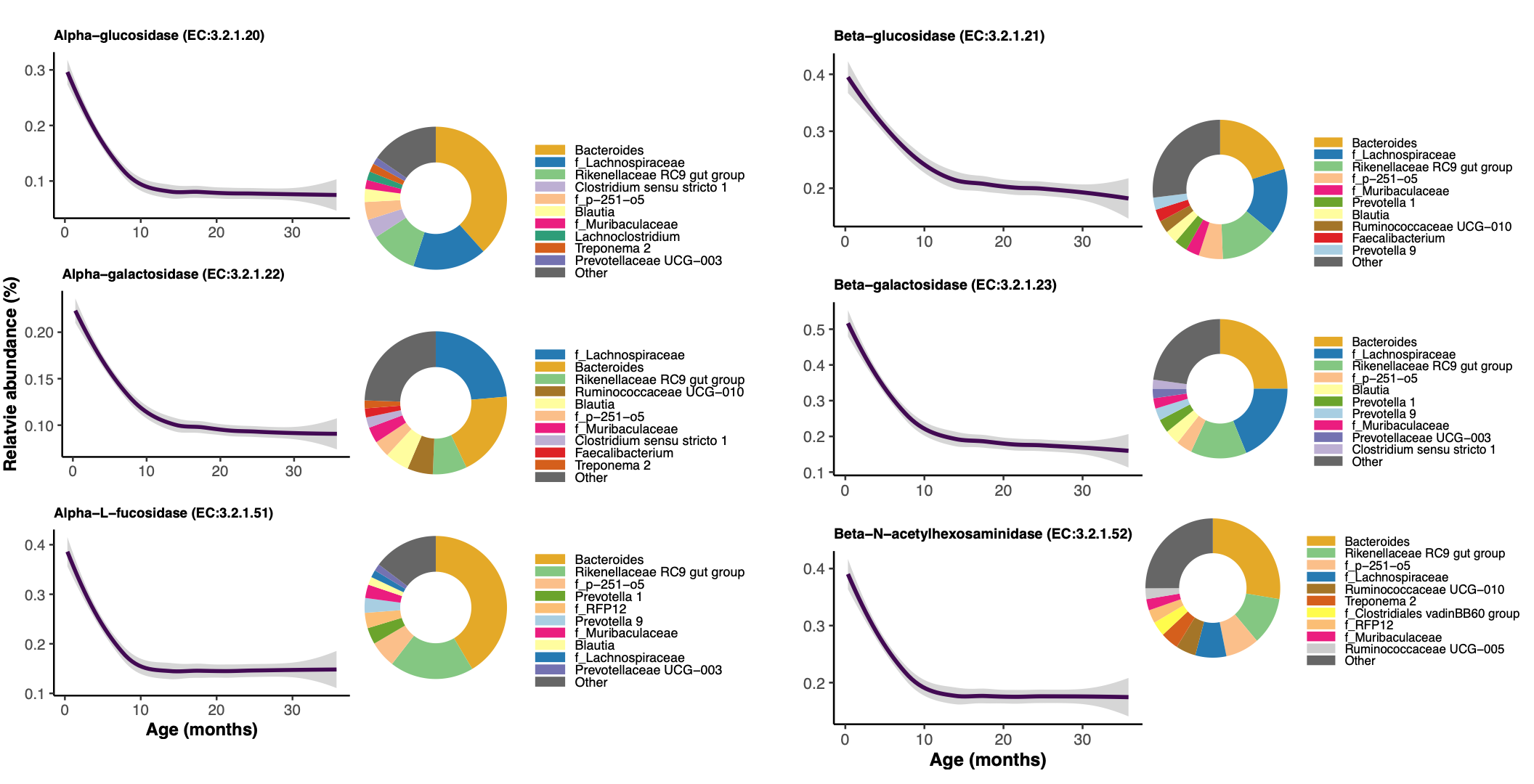

**Figure S9. Age-associated pattern of alpha beta diversity, according to maternal parity.** (A) Alpha diversity was calculated by the Shannon index (richness and evenness of Amplicon Sequencing Variants, ASVs). (B) Beta diversity representation originates from the projection of the first principal component (PC1) of a Principal Component Analysis (PCA) ordination (based on the Aitchison dissimilarity index). The dataset was rarefied at 20,000 reads for the figure.

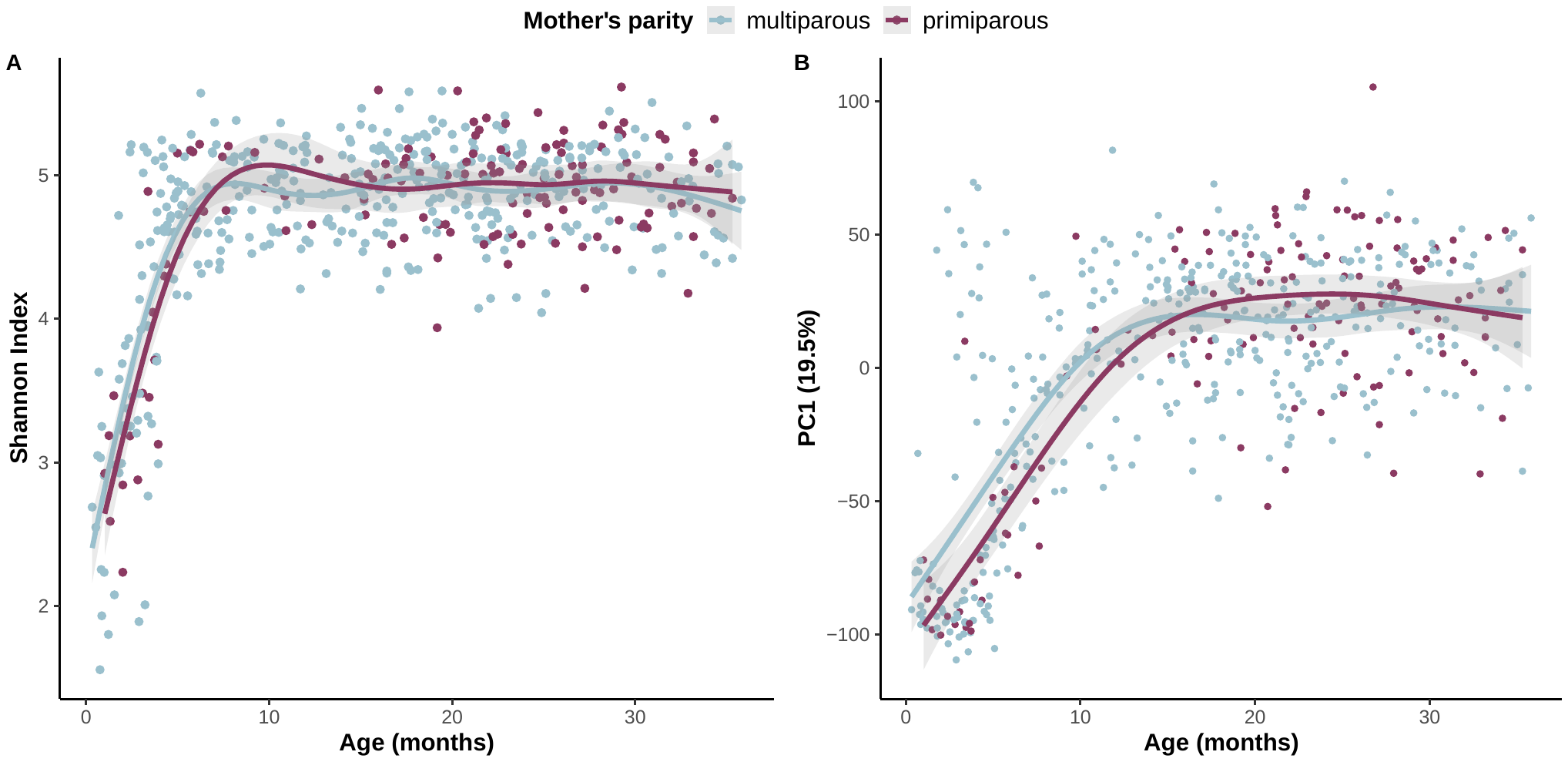

**Figure S10.** Predicted enzymes (EC numbers) that are more abundant in infants (<12 months) born to primiparous females than multiparous females.

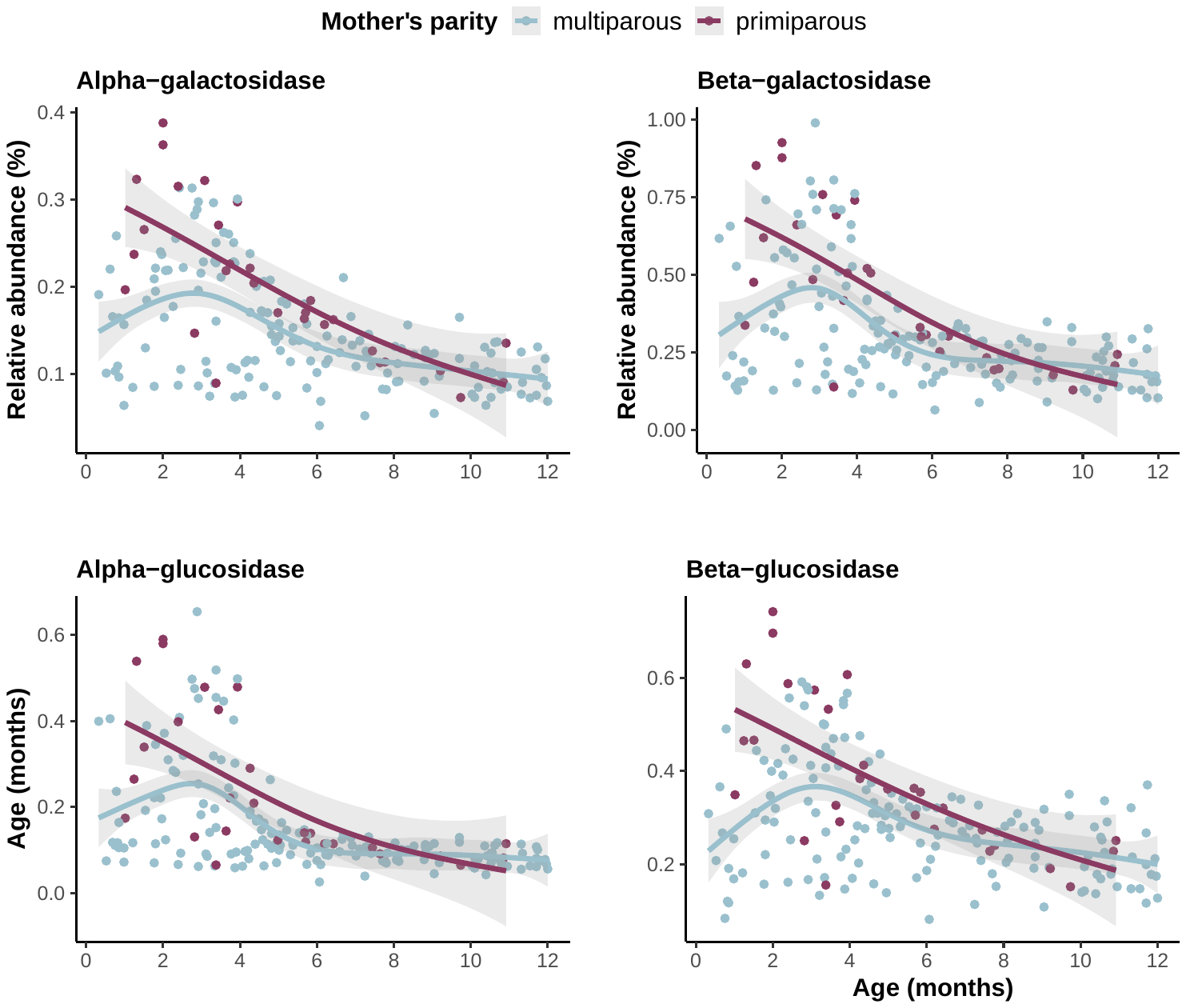

**Figure S11.** Relative abundance of bacterial (A) families and (B) genera that tend to be more abundant in young infants (<12 months) born to primiparous females than multiparous females. These bacteria are all involved in milk sugar degradation.

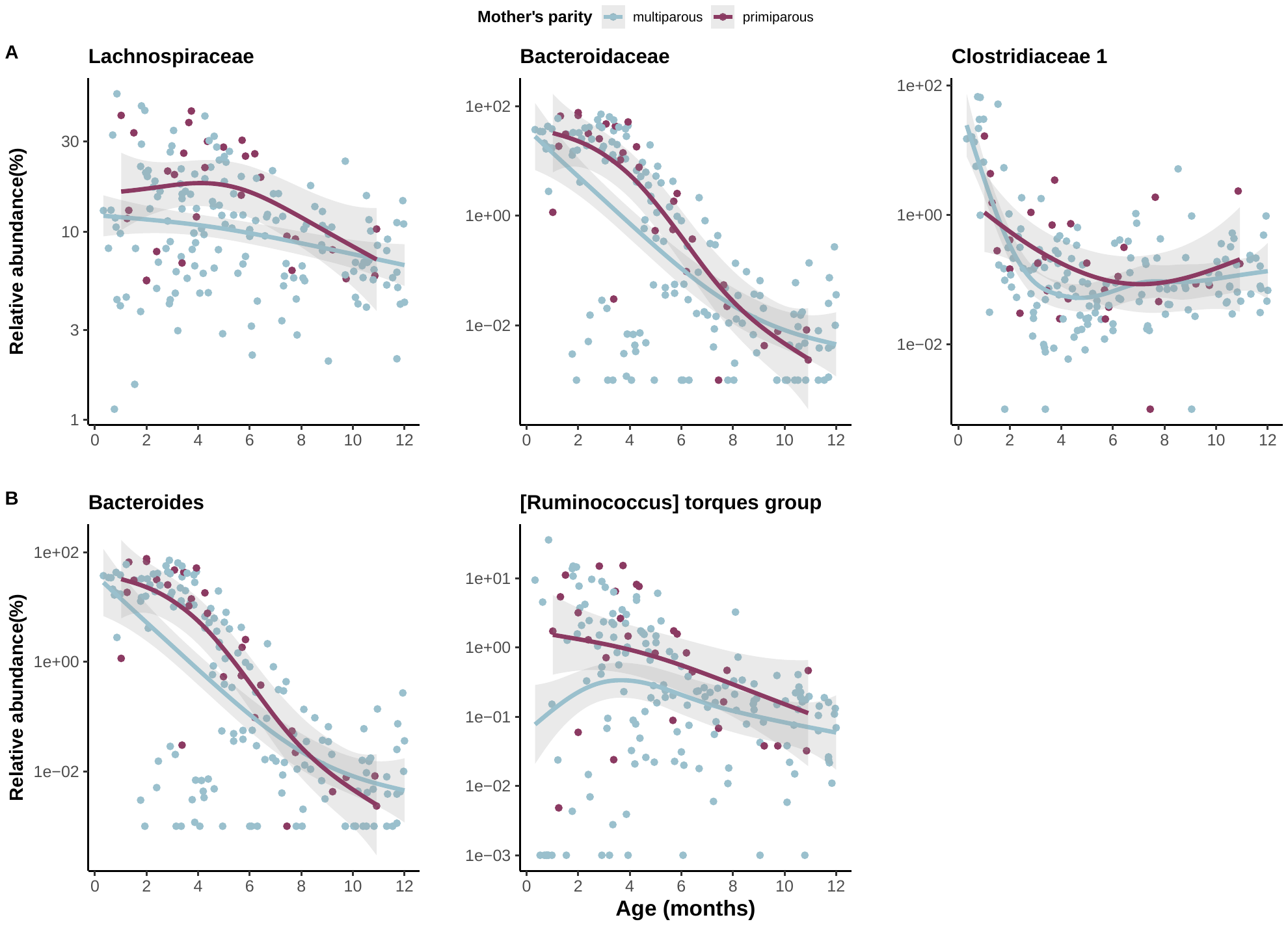

**Figure S12.** **Mother-to-infant transmission of gut microbiome.** Results of the nonparametric resampling approach testing if offspring have a more similar gut microbiome composition (weighted UniFrac dissimilarity) to their mother than to random adult females of the population. The histograms show the random distribution of beta diversity (i.e., when matching each infant sample to a random female sample collected during the same season with 1000 repetitions). The vertical line shows the observed value of beta diversity (i.e. between the actual mother-offspring pairs of fecal samples collected the same day). This analysis was performed separately for young (nursing) infants (aged between 0-12 months, N=136 samples) and old immatures (>18 months, N=201).

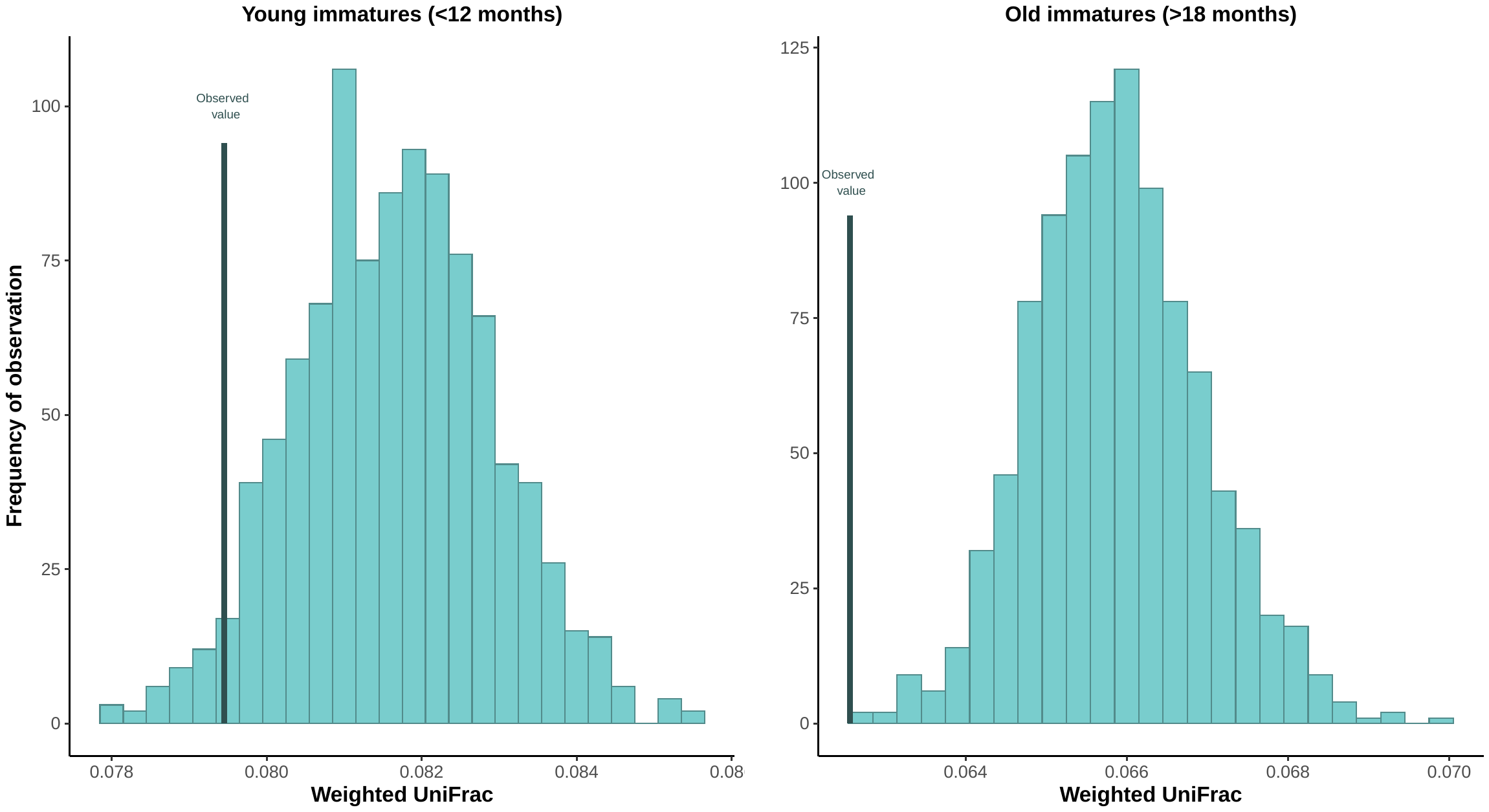

**Legend of Supplementary Tables**

**Table S1. Predictors of alpha diversity in immatures.** Three complementary metrics of alpha diversity (Shannon Index, Observed richness, Faith’s phylogenetic diversity) were modeled using General Additive Mixed Models (GAMMs). Age was included as a smooth term and other covariates (immature sex, maternal rank and parity, cumulative monthly rainfall, average monthly minimum temperature, and log-transformed sequencing depth) as fixed terms. Individual identity and unit membership were included as random effects (i.e. as smooth terms with bs=”re”). For smooth terms, we report the effective degrees of freedom (edf) indicating how the relationship with the smooth term and the response variable is far from a linear relationship (values close to 1 indicate a linear relationship and above 1 a wiggly relationship), the F test statistic, and the p-values. For fixed terms, we report the estimate, t test statistic, and the p-values. The models were run on all samples (0-36 months, N=525) or only samples from young infants (<12 months, N=184) and only samples from old immatures (>18 months, N=259) separately to compare the strength of maternal effect during the nursing and post-weaning period. P-values <0.05 are highlighted in bold.

**Table S2. Predictors of beta diversity in immatures.** We carried out a PERMANOVA using the Aitchison dissimilarity distance between samples (i.e. the Euclidean distance between samples after centered log-ratio transformation of the raw counts) across 10,000 permutations. Individual identity was included as a blocking factor (“strata”) to control for repeated sampling among individuals. The analyses were run on all samples (0-36 months, N=525) or only samples from young infants (<12 months, N=184) and old immatures (>18 months, N=259) separately. The R-squared values indicate the amount of between-sample variation explained by each variable. P-values <0.05 are highlighted in bold.

**Table S3. Predictors of the similarity in gut microbiome composition between mothers- offspring pairs.** We used GAMMs to model the number of shared ASVs and the beta diversity dissimilarity (unweighted and weighted UniFrac) among the mother-offspring matched samples. Age was included as a smooth term and other covariates (immature sex, maternal rank and parity, cumulative monthly rainfall, average monthly minimum temperature) as fixed terms. Individual identity and unit membership were included as random effects (i.e. as smooth terms with bs=”re”). For smooth terms, we report the effective degrees of freedom (edf) indicating how the relationship with the smooth term and the response variable is far from a linear relationship (values close to 1 indicate a linear relationship and above 1 a wiggly relationship), the F test statistic, and the p-values. For fixed terms, we report the estimate, t test statistic, and the p-values. Samples were rarefied at 20,000 reads to control for differences in sequencing depth between immature and mother samples. The models were run on all samples (0-36 months, N=398) or only samples from young infants (<12 months, N=136) and only samples from old immatures (>18 months, N=201) separately to compare the strength of vertical transmission during the nursing and post-weaning period. P-values <0.05 are highlighted in bold.

**Table S4. Results of the hierarchical clustering of the microbial families and genera.** For each family or genus, an ARIMA model was fitted on the centered log-ratio (clr) transformed count per sample. Families were then grouped into 4 clusters based on similarities in age-associated abundance trajectories. A description of the cluster trajectory is provided.

**Table S5. Results of the hierarchical clustering of the Kyoto Encyclopedia of Genes and Genomes (KEGG) Orthologs (KOs) at level 2 and 3 of the BRITE map.** For each metabolic pathway, an ARIMA model was fitted on the relative abundance count per sample. Pathways were then grouped into 4 clusters based on similarities in age-associated abundance trajectories. A description of the cluster trajectory is provided.

**Table S6. Results of the hierarchical clustering of the Enzyme Commission (EC) numbers.** For each enzymatic pathway, an ARIMA model was fitted on the relative abundance count per sample. Pathways were then grouped into 4 clusters based on similarities in age-associated abundance trajectories. A description of the cluster trajectory is provided.

**Table S7. Loading scores of ASVs on the first principal component (PC1).** A Principal Components Analysis (PCA) on the Aitchison dissimilarity matrix was used to examine how immature samples clustered by age. We extracted the loading scores for each ASV onto the first Principal Component (PC1) of the PCA to determine which specific ASVs have the highest influence on the clustering by age of samples. Negative loadings correspond to ASVs that are typically observed in early life (e.g. <-0.4), while positive loadings correspond to ASVs typically observed in later life (e.g., >0.4) .

**Table S8. Predictors of the relative abundance of the microbial families and genera in nursing infants (0-12 months).** We tested for differential abundance for each microbial taxon by fitting a GAMM on the relative abundance of the taxon per sample (log-transformed). Age was included as a smooth term and other covariates (immature sex, maternal rank and parity, cumulative monthly rainfall, average monthly minimum temperature) as fixed terms. Individual identity and unit membership were included as random effects (i.e. as smooth terms with bs=”re”). For smooth terms, we report the effective degrees of freedom (edf) and the p-values. For fixed terms, we report the estimate and the p-values. P-values were adjusted for multiple hypothesis testing by calculating the Benjamini-Hochberg FDR multiple-test correction. The deviance explained by the model is provided.

**Table S9. Predictors of the relative abundance of KEGG Orthologs (KOs) at level 2 and 3 of the BRITE map in nursing infants (0-12 months).** We examined whether any metabolic pathways were differentially abundant across maternal categories (e.g., primi- vs. multiparous) by fitting GAMM on the relative abundance of each metabolic pathway per sample. Age was included as a smooth term and other covariates (immature sex, maternal rank and parity, cumulative monthly rainfall, average monthly minimum temperature, and sequencing depth) as fixed terms. Individual identity and unit membership were included as random effects (i.e. as smooth terms with bs=”re”). For smooth terms, we report the effective degrees of freedom (edf) and the p-values. For fixed terms, we report the estimate and the p-values. P-values were adjusted for multiple hypothesis testing by calculating the Benjamini-Hochberg FDR multiple-test correction. The deviance explained by the model is provided. The pathways and p-values that are significantly different according to maternal parity are in bold.

**Table S10. Predictors of the relative abundance of the Enzyme Commission (EC) numbers in nursing infants (0-12 months).** We ran differential abundance models only on young infants to examine if maternal effects are stronger earlier in development. We tested for differential abundance in each enzymatic pathway by fitting GAMM on the relative abundance per sample. Age was included as a smooth term and other covariates as fixed terms. Individual identity and unit membership were included as random effects (i.e. as smooth terms with bs=”re”). For smooth terms, we report the effective degrees of freedom (edf) indicating how the relationship with the smooth term and the response variable is far from a linear relationship (values close to 1 indicate a linear relationship and above 1 a wiggly relationship) and the p-values. For fixed terms, we report the estimate and the p-values. The deviance explained by the model is provided. The pathways that are significantly different according to maternal parity are in bold.

**Table S11.** **Results of the nonparametric resampling approach testing if maternal and immature gut microbiome communities are more similar than expected by chance.** We compared the number of shared ASVs and beta diversity dissimilarity (unweighted and weighted UniFrac) between the actual mother-offspring matched fecal samples (the observed value) and between random pairs of fecal samples among the immature and a female of the population (the random distribution, with 1000 repetitions). Because of our limited sample size, the random marching was done by either matching the immature sample to (i) a female of the same unit (to control for higher similarity only due to sharing the same social group) or (ii) a female collected in the same season (to control for higher similarity only due to seasonality). Both analyses provide qualitatively similar results. For each random resampling, a GAMM was used to compare the observed and random distribution of the metrics. The type of pair (observed versus random), immature age (as a smooth term) and immature sex were fit. Infant and female identity were included as random effects. The observed value is the average metric across the actual mother-offspring pairs. The random value is the average metric across the random infant-non-mother pairs (average across the 1000 resampling). We report the exact p-value (calculated as the proportion of models with positive estimates for the number of shared ASVs and the proportion of models with negative estimates for beta dissimilarity) and the 95% confidence interval of the estimate. Fecal samples were rarefied at 20,000 reads to control for difference of sequence depth between immature and female samples. These analyses were run including all immatures samples (0-3 years), only young infant samples (<12 months) or only old immatures (>18 months) to compare the strength of the effect among the different age categories.

**Table 12.** **Summary of the ASVs shared between mother-offspring pairs in early life (<12 months).** The 136 pairs of matched fecal samples between mother and offspring (<12 months) were rarefied at 20,000 reads to control for difference of sequence depth between infant and mother samples. Out of the 3402 ASVs observed in young infants, 1615 were shared by at least one mother-infant pair. For each ASV, we report its relative prevalence (i.e. the proportion of samples in which the ASV is found) and its average relative abundance among the infant samples, as well as the proportion of mother-offspring pairs that share this ASV. The loading score of each ASV onto the first Principal Component (PC1) of the PCA (based on Aitchison dissimilarity matrix between infant samples) is provided. The ASVs with the most negative loading scores (<-0.4) characteristic of early life are not frequently shared between mother-offspring pairs.

**Table S13. Predictors of beta diversity in immatures.** We carried out a PERMANOVA using unweighted and weighted UniFrac dissimilarity distance between samples (on the raw counts) and 10,000 permutations. Individual identity was included as a blocking factor (“strata”) to control for repeated sampling among individuals. The analyses were run on all samples (0-36 months, N=525) or only samples from young infants (<12 months, N=184) and old immatures (>18 months, N=259) separately. The R-squared values indicate the amount of between-sample variation explained by each variable. P-values <0.05 are highlighted in bold.
